# Long-term perceptual priors drive confidence bias that favors prior-congruent evidence

**DOI:** 10.1101/2024.06.17.599305

**Authors:** Marika Constant, Elisa Filevich, Pascal Mamassian

## Abstract

According to the Bayesian framework, both our perceptual decisions and confidence about those decisions are based on the precision-weighted integration of prior expectations and incoming sensory information. While it is generally assumed that priors influence both decisions and confidence in the same way, previous work has found priors to have a stronger impact at the confidence level, challenging this assumption. However, these patterns were found for high-level probabilistic expectations that are flexibly induced in the task context. It remains unclear whether this generalizes to low-level perceptual priors that are naturally formed through long term exposure. Here we investigated human participants’ confidence in decisions made under the influence of a long-term perceptual prior: the slow-motion prior. Participants viewed tilted moving-line stimuli for which the slow-motion prior biases the perceived motion direction. On each trial, they made two consecutive motion direction decisions followed by a confidence decision. We contrasted two conditions – one in which the prior impacted discrimination performance, and one in which it did not. We found a confidence bias favoring the condition in which the prior influenced discrimination decisions, even after accounting for performance differences. Computational modeling revealed this effect to be best explained by confidence using the prior-congruent evidence as an additional cue, beyond the posterior evidence used in the perceptual decision. This is in agreement with a confirmatory confidence bias favoring evidence congruent with low-level perceptual priors, revealing that, in line with high-level expectations, even long-term priors have a greater influence on the metacognitive level than on perceptual decisions.

**Author Summary:** Prior expectations play a critical role in shaping not only the perceptual inferences that we make, but also how confident we feel about those inferences. Bayesian confidence models capture that role, but assume priors to influence both decisions and confidence in the same way. Against this assumption, previous work has found dissociations in the influence of priors on decisions and confidence. However, that work has focussed only on high-level probabilistic priors, rather than the low-level perceptual priors that constrain our processing across many naturalistic situations. Here, we examine whether such dissociations arise under the influence of a low-level perceptual prior that naturally affects humans’ perception of motion, namely, the expectation that objects move slowly. We reveal evidence for such a dissociation: prior-congruent evidence impacts confidence to a greater extent than perceptual decisions. This suggests the existence of an implicit confidence bias favoring information that confirms prior beliefs, even in the case of long-term perceptual priors.

---

Our perception of the environment is constantly compromised by uncertainties, and the Bayesian framework has become a popular way to account for how we cope with this uncertainty across a variety of cases (1–4). According to Bayesian inference models of perception, our perceptual experience and decisions depend not only on incoming sensory information (the “likelihood”), but also on prior expectations (“priors”). These sources of evidence are combined with weights proportional to their precision to form the “posterior” probability distribution function, and the perceptual judgment is obtained by applying a decision rule on this posterior probability. More recently, these Bayesian decision models have also been extended to account for our confidence judgments about the validity of our perceptual decisions, with confidence captured as the estimated posterior probability of being correct (5–10). This implies a role for priors in confidence as well, which has been supported empirically in both humans (11–13) and non-human animals (14).

Recent evidence suggests that priors may have dissociable impacts on perceptual decisions (first-order) and confidence judgments about those decisions (second-order). For example, priors about sensory precision, performance, or task difficulty have been shown to affect second-order judgments without changing first-order performance (15–17). Arguably, however, this does not constitute strong evidence for a differential impact of prior information on first- and second-order decisions: These higher-order priors concerned specifically the confidence-related features of the signal, but were irrelevant for perceptual decisions. Clearer evidence comes from recent work examining how decision-relevant priors are weighted in confidence relative to their use in discrimination decisions. This work found robust dissociations between these processing levels: Prior information was used to a greater extent in subjective confidence than in the decisions themselves (18). This suggests that priors have a particular role in confidence, over and above their effect on perceptual decisions. However, that work – along with most of the work on priors in confidence aside from a recent counter-example (19) –, manipulated priors based on probabilistic expectations that were flexibly induced in the task context and likely acted post-perceptually. One plausible explanation for these ‘high-level’ priors especially influencing confidence is that the abstract probabilistic information that they carry is difficult to integrate with perceptual information, but easier to integrate with confidence information. Additionally, while it may be advantageous not to rely on priors for perceptual decisions when they flexibly and rapidly change, this would not be the case for more stable, slower-updating priors. In order to test these explanations, a natural step is to ask whether the same asymmetries follow from priors that are neither difficult to integrate perceptually, nor rapidly changing, as is the case for low-level perceptual priors.

Low-level priors are often formed naturally across a breadth of life experience, such as a prior to perceive light as coming from above (20), and are sometimes therefore referred to as ‘long-term’ priors. Further, they have been suggested to act differently to high-level expectations in a variety of other ways, being slower to update, more cognitively impenetrable, and more context-independent (21–23). Some have also suggested long-term priors to be implemented differently, in a more bottom-up processing stream (23,24), although there is recent counter-evidence to this argument (25). In light of these proposed differences, it remains unclear whether the same dissociations between decisions and confidence that emerge under the influence of high-level expectations would hold for long-term perceptual priors. Here, we test this question by investigating confidence under the influence of a long-term, naturally formed prior. To do so, we focus on the ‘slow-motion’ prior.

It has been argued that humans have a low-level perceptual prior for motion speed to be slow, which is thought to originate from the fact that many objects we perceive in the world are stationary or move very little (26). This has been shown to impact motion perception across different tasks and to explain a variety of biases in motion perception (26–29). In line with Bayesian theory, the slow-motion prior influences inferences about motion particularly when sensory information is uncertain, or in other words when likelihoods are noisy. One example of uncertain motion information occurs when lines whose ends are poorly visible move through an aperture, producing the so-called “aperture effect”. Under these conditions, the direction and speed of the line motion are inherently ambiguous and participants are initially biased to perceive a motion direction that is exactly orthogonal to the line orientation (30,31). This is consistent with perceiving the slowest possible motion that can still explain the sensory information, in line with the idea that the slow-motion prior influences perceptual inference (26,32). Sotiropoulos and colleagues found that this bias towards orthogonal motion emerged strongly in participants’ motion direction decisions and was well captured by a Bayesian decision model including the slow-motion prior (32). That work also found that the bias was reduced and even reversed by exposing participants to fast motion between test sessions, further demonstrating the impact of motion priors in this perceptual effect. Here, we build on similar stimuli as those used by Sotiropoulos et al. to investigate the effects of the slow-motion prior on perceptual decisions and confidence.

We measured confidence sensitivity and bias with the confidence forced-choice paradigm (33). On every trial in this paradigm, participants made two consecutive perceptual decisions followed by a confidence choice regarding which of the two decisions was more likely to be correct. Each perceptual decision consisted in reporting whether a set of lines was perceived to translate in a direction that was clockwise or counterclockwise relative to a reference. We contrasted two conditions in order to examine the impact of information from the slow-motion prior on perceptual decisions and confidence judgments. One condition was tailored so that the slow-motion prior contributed strongly to whether the perceived motion direction was expected to be clockwise or counterclockwise relative to the reference, and the other condition was such that the prior was uninformative to the motion direction decision. By matching the first-order choice rates across these conditions, we could then examine whether there was a residual confidence bias favoring one of the conditions. If so, this could suggest that confidence uses the prior information differently than the first-order decisions do. Further, to quantify this effect and better understand the mechanisms underlying the use of perceptual priors in confidence, we fit and compared several models of confidence choices under the influence of priors.

## Results

On every trial of the confidence forced-choice task, participants decided about the motion direction of a set of parallel lines moving through a circular aperture (Fig. 1A) on two consecutive and independent stimulus intervals. The preferred motion direction due to the slow-motion prior was orthogonal to the orientation of the moving lines. On the outside of the circular aperture, two abutting arcs, each subtending 90°, served as choice regions – one orange, one blue – and defining the ‘reference’ at their separation (Fig. 1A). In each interval participants made a decision about whether the motion direction was towards the displayed blue or orange region. Participants then made a confidence forced-choice on every trial about whether they were more confident about being correct in the first or second interval (Fig. 1B), thereby removing issues of confidence biases that come with subjective ratings (34). We created two conditions. In the Bias condition, the preferred direction – orthogonal to the line orientation – fell within either the orange or blue region, thus creating a decision bias (Fig. 1A). In the No-Bias condition, the preferred motion direction(s) fell exactly 90° away from the reference and hence there was no bias favoring either decision. In this way, we could compare the No-Bias condition, in which decision performance was determined entirely by the incoming sensory information (the likelihood), against the Bias condition, in which the slow-motion prior influenced performance. The strength of the likelihood, or the difficulty of the stimuli, was controlled by the angle, θ, subtended by the true motion direction and the reference.

**Figure 1.**
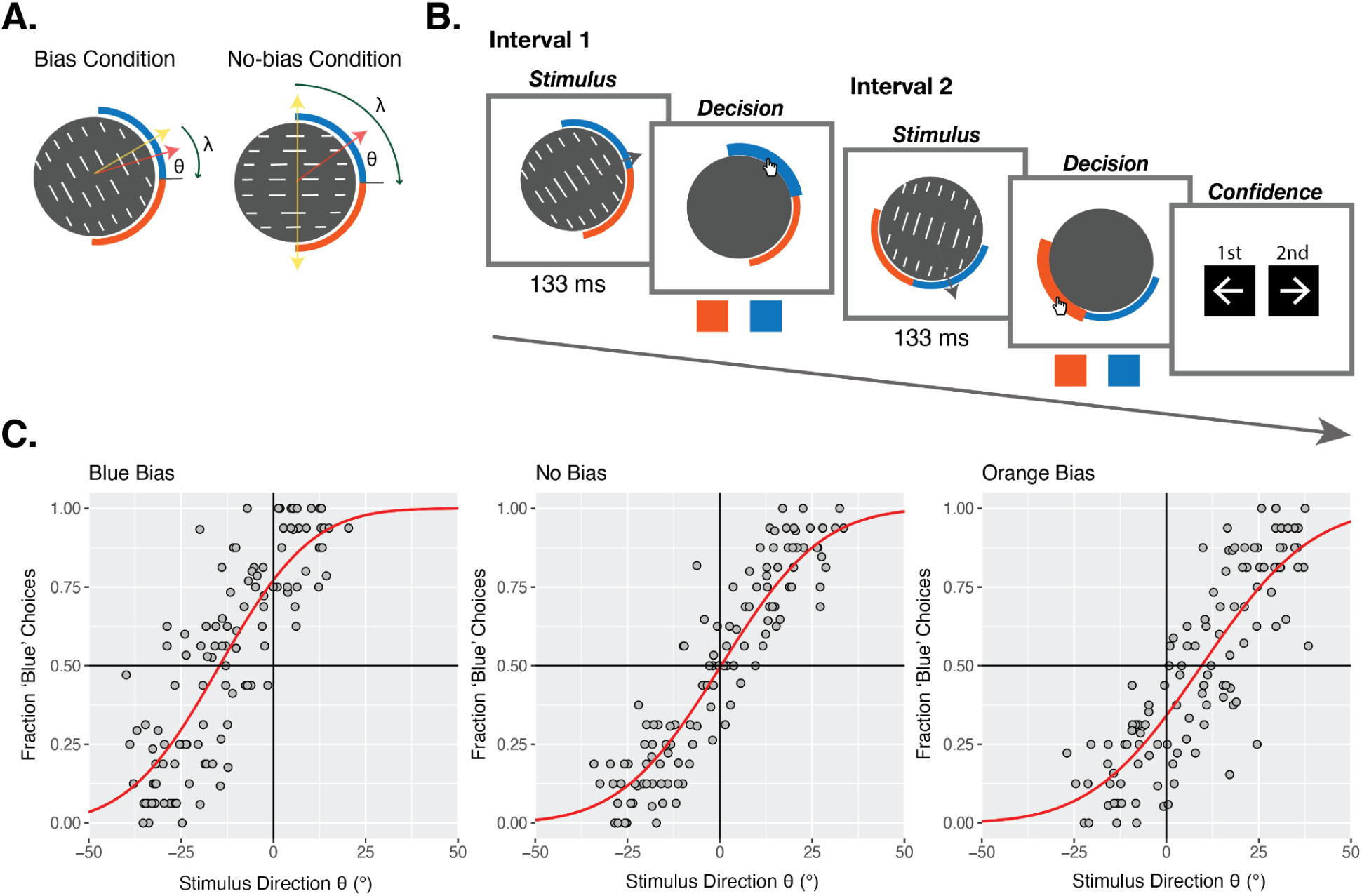
Task and Manipulations. **A. Conditions and sketch of stimuli.** The stimuli consisted of a set of lines moving through a circular aperture, shown at low contrast and for a short duration. The yellow arrows indicate the preferred motion directions consistent with the prior, orthogonal to the lines’ orientation. In the Bias condition, the angle, λ, between this prior and the reference was 35°, creating a decision bias favoring the color in which the prior falls. In the No-Bias condition, λ was 90°, such that we expected no prior-induced decision bias favoring either color. The red arrows indicate the true motion direction on an example stimulus. The strength of the likelihood was controlled by the angle, θ, between this motion direction and the reference. A larger θ was required in the No-Bias condition to reach the same choice rates of seeing a motion direction in the blue region, since the prior favored blue and orange decisions equally. **B. Experimental task.** A trial of the confidence forced-choice task included two stimulus intervals, in which participants saw a moving stimulus and made an orange/blue decision about the motion direction of the lines, followed by a confidence forced-choice about which of their responses they were more confident of being correct. **C. Manipulation check.** Pooled results from the adaptive staircase (ASA) procedure in which many different θ values were tested in their effect on first-order decisions in both conditions before the full confidence task. Each gray point corresponds to the responses for a given stimulus intensity level from one participant. θ values towards the blue region are encoded as positive, and towards the orange region are encoded as negative. Red psychometric functions show the fit cumulative normal distribution function capturing the relationship between the θ and the fraction of blue choices. In the No-Bias condition (middle panel), there is no choice bias for orange or blue. In the Bias condition, when the prior falls in the blue region (left panel), the function is shifted to the left, showing a successfully induced choice bias favoring blue. When the prior falls in the orange region (right panel), the function is shifted to the right, showing a successfully induced choice bias favoring orange.

### Manipulation Check

We predicted that the slow-motion prior would bias participants’ decisions in the Bias condition, and hence that the Bias condition would require a smaller θ angle in order to lead to the same decision rates as the No-Bias condition. This effect would serve as a basic manipulation check to ensure that the conditions and slow-motion prior had the intended impact. Additionally, it was necessary to quantify this effect in each participant before the experiment such that we could choose the θ values that would be expected to lead to matched decision rates. To do this, before the main task we ran a staircasing procedure which sampled different θ values in both conditions and then fit psychometric functions to these data. The fitted psychometric functions capturing the bias effect across all pooled participants are shown in Fig. 1C, and this bias effect was verified in each individual participant (Fig. S1). Motion towards the orange region was encoded with negative θ’s and blue region with positive θ’s. We found a clear bias effect, such that the psychometric was shifted leftward when the preferred motion direction from the prior was in the blue region and rightward when it was in the orange region, compared to the No-Bias condition (Fig. 1C). The point of subjective equality (PSE) when there was a blue bias was −14.57°, meaning that when the line motion direction was 14.57° into the orange region, participants nonetheless chose orange and blue at equal rates. In contrast, the PSE when there was an orange bias was 9.53°, and the PSE in the No-Bias condition was 0.30°. Together, this suggests that the bias manipulation worked as planned.

### Motion-Direction Decisions

After running the staircasing procedure, we chose the values of θ angles that corresponded to a 0.15, 0.35, 0.65 and 0.85 expected probability of choosing blue (P(‘Blue’)) for each participant. This procedure was followed both for the No-Bias condition and for the Bias condition (for a total of eight different θ values). In the Bias condition, the values corresponding to a 0.15 and 0.35 P(‘Blue’) were taken from cases in which the slow-motion prior induced an orange bias (the lines were oriented to have the orthogonal direction in the orange region), and values corresponding to a 0.65 and 0.85 P(‘Blue’) were taken from cases in which the prior induced a blue bias (the lines were oriented to have the orthogonal direction in the blue region). Overall, this meant that we chose stimulus levels that were expected to lead to matched choice probabilities between conditions. Crucially, however, the choice rates in the Bias condition would be due in part to the slow-motion prior; whereas the choice rates in the No-Bias condition would be solely driven by the stimulus evidence. These four θ values in each condition were paired in all possible combinations for the two intervals of the confidence forced-choice task, except for pairs repeating the identical θ in the same condition.

To check that the choice rates were matched overall across the two conditions, we ran a logistic mixed effects model on Response including Expected P(‘Blue’) (as an index of the stimulus intensity setting), Condition (Bias vs No-Bias), and their interaction as fixed effects, as well as by-participant random intercepts. If the chosen θ values from the staircasing procedure failed to produce matched choice rates between conditions, this will generate an interaction between Expected P(‘Blue’) and Condition. Indeed, a significant interaction, *χ*^2^(3)=302.97, p<0.001, BF_10_ = 1.19 × 10^60^, revealed that the response rate per stimulus setting additionally depended on the condition, making our analysis more complex. The response rates were more extreme in the Bias condition (Fig. 2A), with higher odds of choosing blue when the stimulus was blue (Expected P(‘Blue’) of 0.65: odds ratio of the Bias to the No-Bias condition (OR_Bias/No-Bias_) = 1.55, 95% confidence interval (CI) [1.32, 1.81], Z=8.75, p<0.001; Expected P(‘Blue’) of 0.85: OR_Bias/No-Bias_ = 1.13, 95% CI [0.92, 1.38], Z=1.81, p=0.071), and lower odds when the stimulus was orange (Expected P(‘Blue’) of 0.15: OR_Bias/No-Bias_ = 0.54, 95% CI [0.42, 0.69], Z=-7.91, p<0.001; Expected P(‘Blue’) of 0.35: OR_Bias/No-Bias_ = 0.49, 95% CI [0.41, 0.58], Z=-13.43, p<0.001), compared to the No-Bias condition, although this contrast was not significant for the strongest blue setting. The more extreme response rates in the Bias condition suggest that the bias effect was stronger during the main experimental trials, compared to what was calibrated in the staircasing procedure. To quantify this, we estimated the strength of the bias effect in the experiment by fitting a cumulative normal distribution function with one symmetrical bias level (+/- μ_Bias_) for the orange and blue directions relative to the No-Bias condition, which we fit separately as a baseline (Fig. 2A). The sensory sensitivity was also allowed to differ between conditions. This was done for each participant and the resulting mean μ_Bias_ across participants was 20.81° (Fig. 2A), indeed indicating a stronger bias effect than in the staircasing (Fig. 1C). Hence, we needed to account for this residual difference in first-order response rates in our analysis when testing for a confidence bias.

**Figure 2.**
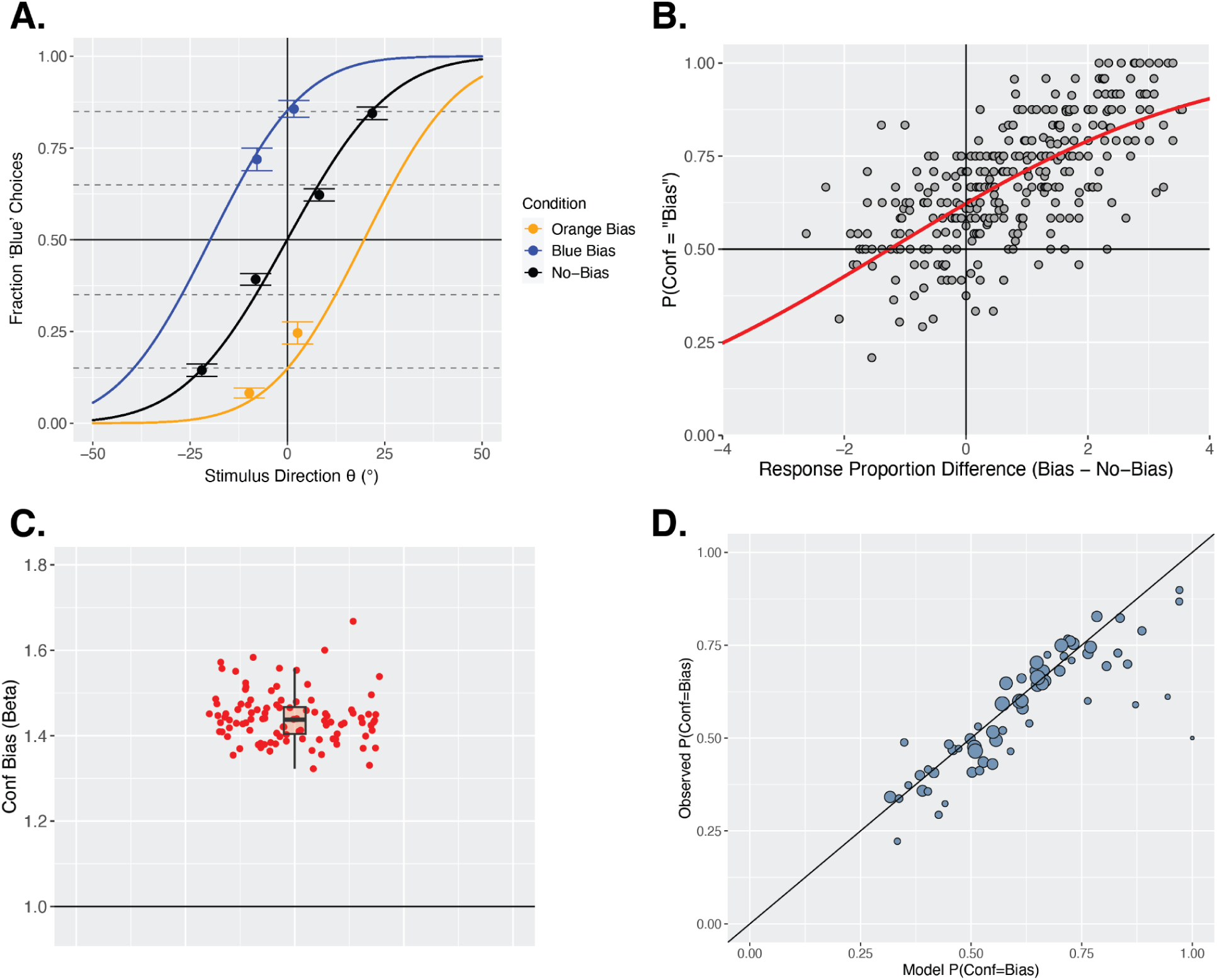
Motion-Direction and Confidence Decision Results. **A. Motion-direction decisions.** Choice results for the first-order orange/blue decisions in each condition. The four intensity levels were targeting 0.15, 0.35, 0.65 and 0.85 probabilities of choosing blue (P(‘Blue’)) in each condition. In the Bias condition, this consists of intensity levels targeting 0.65 and 0.85 P(‘Blue’) when there was a blue bias (blue points) and targeting 0.15 and 0.35 P(‘Blue’) when there was an orange bias (orange points). The targeted choice rates are shown with horizontal dashed gray lines. Although we targeted these matched choice rates between conditions, the observed fractions of blue choices were less extreme in the No-Bias condition than in the Bias condition. Error bars reflect the SEM across participants. The psychometric functions were fitted to each individual participant to capture the strength of the bias from the slow-motion prior relative to the No-Bias baseline, and to capture possible differences in sensitivity between conditions. The functions shown here reflect the mean fit bias (μ) and noise (σ) values across participants. The mean μ_Bias_ was 20.81, mean μ_No-Bias_ was 0.44, mean σ_Bias_ was 18.50, and mean σ_No-Bias_ was 20.46. **B. Non-parametric confidence bias results.** On the x axis is the difference in the z-scored rate of choosing the expected color between the Bias and No-Bias condition. Each gray point represents this difference for one pair of stimulus strengths assigned to the two intervals of a confidence pair (16 points per participant). On the y axis is the probability of choosing the Bias condition interval as the more confident interval. When there is no difference in response proportions between conditions (x=0), and the perceptual difficulties are therefore matched for that pair of settings, we expect equal confidence choices between conditions. If this is not the case, this indicates a confidence bias. The red psychometric function captures the fit cumulative normal distribution function to the relationship between this response proportion difference and the confidence choice rates from the pooled data. The PSE is shifted leftward to −1.26 (SEM=0.44 from performing this analysis in individual participants, see Fig. S2), indicating a confidence bias favoring the Bias condition. **C. CFC-model confidence bias results.** We fit the *cfc-model* (33) across 100 bootstrapped runs and extracted the fit confidence bias parameter *β* from each. Each red point corresponds to one fit *β* parameter result. The boxplot shows the median, interquartile range (IQR) with hinges showing the first and third quartiles, and vertical whiskers stretching to most extreme data point within 1.5*IQR from the hinges. The horizontal black line at y=1 shows the expected *β* if there was no confidence bias. **D. Quality of CFC-model fit.** Predicted against observed confidence choice rates across all participants. Each point shows a pair of stimulus settings with one interval in each condition and a given set of two orange/blue responses, and the size of the point reflects the number of trials. The closer the points are to the x=y line, the better the model is at modeling confidence choice rates.

### Confidence Forced-Choices

We sought to explore whether, after accounting for any residual difference in orange/blue decision rates, there is still a confidence bias favoring one of the conditions. If so, this could indicate a different influence of the prior on perception and confidence, which could then be further explored using Bayesian confidence modeling. Because the orange/blue decision rates were not perfectly matched across conditions, and following our pre-registered plan for this potential situation, we used a non-parametric approach to assess confidence bias, which accounted for potential mismatches in first-order decision rates. We took the difference in z-scored response rates for the preferred stimulus between conditions for every stimulus setting pair for every participant, excluding pairs with both intervals in the same condition, following previous work (35). For example, for the stimulus settings expected to lead to a 0.85 probability of choosing blue in each condition, we took the observed rates of choosing blue, z-scored them to transform them to the (-∞,∞) domain, and took the difference between them. A difference of zero then indicated matched first-order choice rates, positive differences indicated more extreme choice rates in the Bias condition, and negative differences indicated more extreme choice rates in the No-Bias condition. Then, regardless of whether these values were shifted positively overall, as we expect due to the result of more extreme choice rates in the Bias condition, we can fit a cumulative normal distribution function to the relationship between these z-scored response proportion differences and the probability of choosing the Bias condition as the more confident interval. If the PSE is at 0, this indicates that when participants have matched first-order choice rates they also have matched confidence choice rates between conditions, suggesting no confidence bias. If, however, the PSE is shifted away from 0, this indicates a confidence bias. We found a clear negative shift of this function, fit to the pooled participants, with a PSE of −1.26 (SEM=0.44 from performing this analysis in individual participants, see Supplementary Fig. S2), suggesting a confidence bias favoring the Bias condition (Fig. 2B). When response rates are matched (z-scored response proportion difference = 0), the fit psychometric predicts the participants to still choose the Bias condition as the more confident interval at a rate of 0.62.

### Quantifying Confidence Bias: CFC-Model

To further quantify this confidence bias, we fit the *cfc-model* developed in previous work (33) for use with the confidence forced-choice paradigm. This model includes a confidence bias term, *β*, which, for a single task or condition, scales the estimated sensory sensitivity of a participant, such that *β*=1 captures correctly estimated sensitivity, *β*>1 captures an overestimation of sensitivity – indicating overconfidence –, and *β*<1 captures an underestimation of sensitivity – indicating underconfidence. With two different conditions, the model fits *β* as the ratio of biases between them (one can only estimate whether participants are over- or under-confident in one condition relative to the other). In our case, we choose the ‘No-Bias’ condition as baseline, so a fit *β* larger than 1 would indicate an overconfidence in the ‘Bias’ condition. We fit the model to the pooled normalized data from all participants across 100 bootstrapped runs, which revealed the mean fit *β* to be 1.44 (SEM=0.006; Fig. 2C). The quality of the model fit is shown in Fig. 2D. These results, in agreement with the non-parametric analysis above, indicate a confidence bias favoring the Bias condition. While this model can be used to naively quantify a confidence bias relative to first-order choice rates, it does not provide an account of the computations occurring in our design, under the influence of the slow-motion prior. To investigate these computations, in exploratory analyses we consider a Bayesian decision and confidence model, along with several alternative generative models.

### Bayesian Optimal Observer Model

In the Bayesian Optimal Observer model, we first need to define the prior to represent the preferred motion directions that are orthogonal to the lines’ orientation. Even though we relied on the slow-speed prior to bias one condition, this prior has a direct impact on perceived direction (30), and since we are measuring perceived direction rather than speed, we directly model the prior in terms of the preferred motion directions. This motion direction prior is implemented as the sum of two Gaussians of equal variance, whose means are separated by 180° (Fig. 3A). In the No-Bias condition, these means are 90° away from the reference in either direction. In the Bias condition, one mean is 35° away from the reference in the bias direction, and the other is 145° away from the reference in the opposite direction. This latter component is outside the allowed decision region and thus will have a negligible effect in our model (see below). The likelihood is modelled as a single Gaussian centered around the internal signal generated from the stimulus with added internal noise. These combine to form the posterior distribution (Fig. 3A). Orange/blue perceptual decisions in this model are based on the area under the posterior on either side of the sensory reference within the allowed decision region (from −90° to +90°). When this area is larger on the negative side, an “orange” choice is made, and when it is larger on the positive side, a “blue” choice is made (Fig. 3A-B). Following the literature capturing Bayesian confidence as the estimated posterior probability of being correct (5–10), confidence in the decision is then equal to the proportion of the total area (within the allowed decision region) that falls on the chosen side (thus varying between 0.5 and 1). The confidence forced-choice is based on whichever of these confidence values is larger across two intervals. The free parameters of this Bayesian model are the standard deviation of each of the two Gaussians that form the Gaussian mixture prior, σ_P_, and the standard deviation of the likelihood, σ_L_, that represents the sensory uncertainty. These were fit individually for each participant to find the parameters that could best explain their orange/blue decisions. We used a simplification to deal with computational complexity and considered the prior in the Bias condition to be a single Gaussian centered around 35° towards the bias direction. Since most of the other Gaussian fell well outside the allowed decision region, this component had a negligible impact. The results from these model fits are shown in Fig. 3C by simulating from the full unsimplified model, showing that the fit σ_P_ and σ_L_ parameters can still well account for motion-direction decisions.

**Figure 3.**
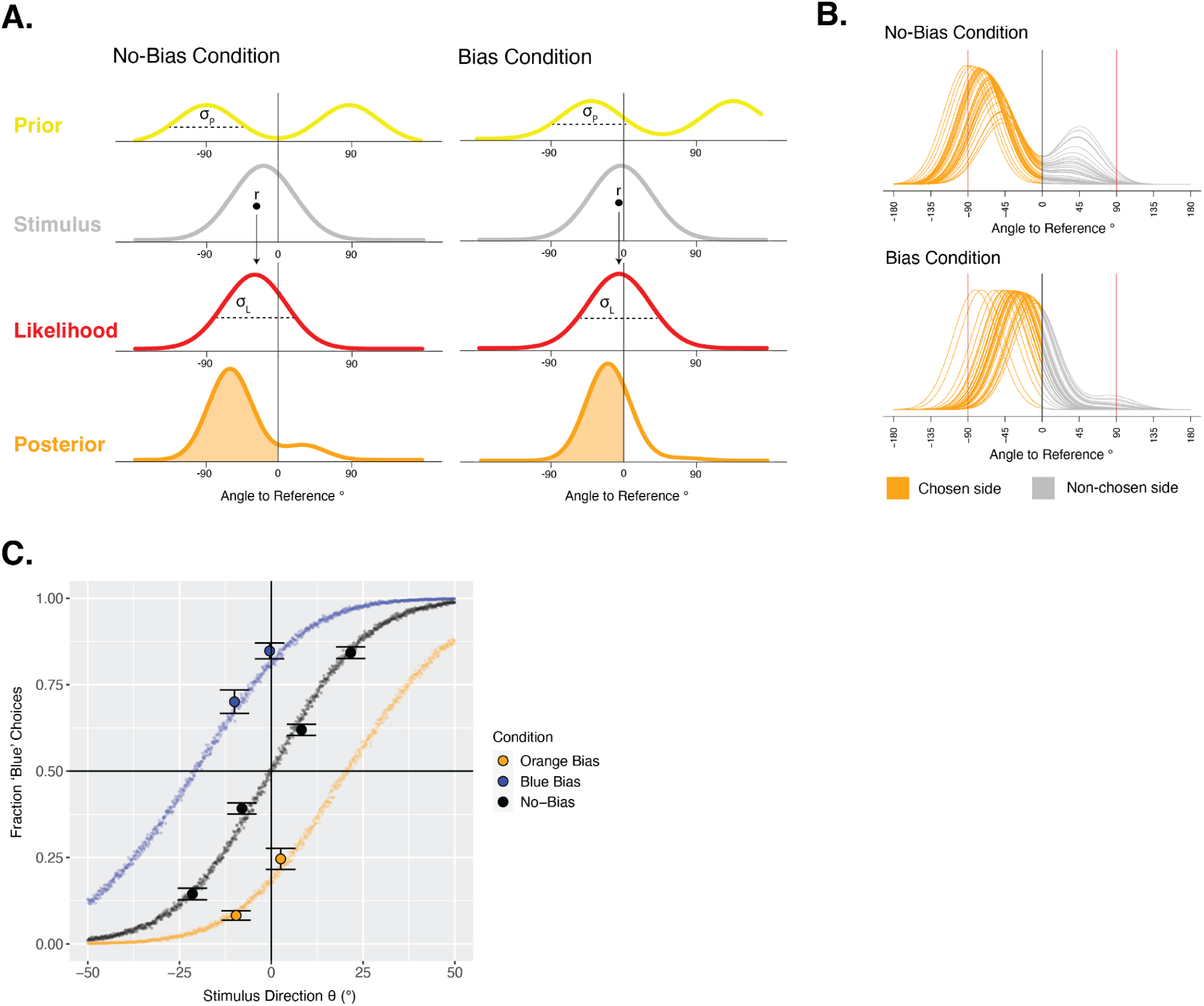
Bayesian Optimal Observer Model. **A. Sketch of Bayesian Optimal Observer model.** The prior on perceived motion direction is a sum of two Gaussians of equal standard deviation (σ_P_), with means at the orthogonal motion directions (90° from the reference in the No-Bias condition and 35°/145° from the reference in the Bias condition). The likelihood comes from the internal signal, *r*, sampled from the stimulus with added internal noise (σ_L_). The prior and likelihood are combined to form the posterior distribution. The shaded orange area under the posterior leads to the first-order decision and confidence. **B. Posteriors per condition from Bayesian Optimal Observer model.** Posterior distributions from the Bayesian Optimal Observer model across 50 samples of an internal signal from the stimulus to form the likelihood, only including samples leading to an orange decision. These reflect posteriors across different trials of the same stimulus intensity and condition, which led to the same response. The confidence in these trials is captured by the ratio of area under the orange part of the posterior (chosen side) to the area under the gray part of the posterior (non-chosen side), between the red vertical lines (allowed choice region). The prior tends to pull the maximum of the posterior in the No-Bias condition away from the reference towards −90°, so the ratio of area under the orange versus gray parts of those posteriors tends to be higher, suggesting higher average confidence in the No-Bias condition despite matched choice rates. **C. Modeled motion-direction decision results.** Results from the fit Bayesian Optimal Observer decision model, simulated across 1001 stimulus intensities ranging continuously (in steps of 0.1) from −50 to 50 using the best-fitting σ_P_ and σ_L_ parameters for each participant, against the data.

We then used this fit model and investigated the confidence forced-choice patterns that would be expected of a Bayesian optimal observer given the first-order choice rates that we found, and compared them to the observed confidence patterns. The raw confidence forced-choice rates are shown in Fig. 4A. Because of the confidence forced-choice task structure and the fact that confidence refers to an estimate that a perceptual decision is correct, we should distinguish confidence choices depending on the stimulus intensity (leading to easy and hard perceptual decisions) and response (blue and orange) of each of two intervals on a given trial. For example, if the Bias condition interval is easy and the No-Bias condition interval is difficult, there is a high probability of a confidence choice favoring the Bias condition. Likewise, if the Bias condition response is correct and the No-Bias condition response is incorrect, there would be a tendency for the confidence choice to favor the Bias condition with all else equal. Hence, the confidence results in Fig. 4A are sorted first into the pairs of responses (forming the four separate panels), and then each grid cell captures a particular pair of stimulus intensities across the two intervals. The overall prominence of red demonstrates the higher rate of confidence choices favoring the Bias condition observed.

**Figure 4.**
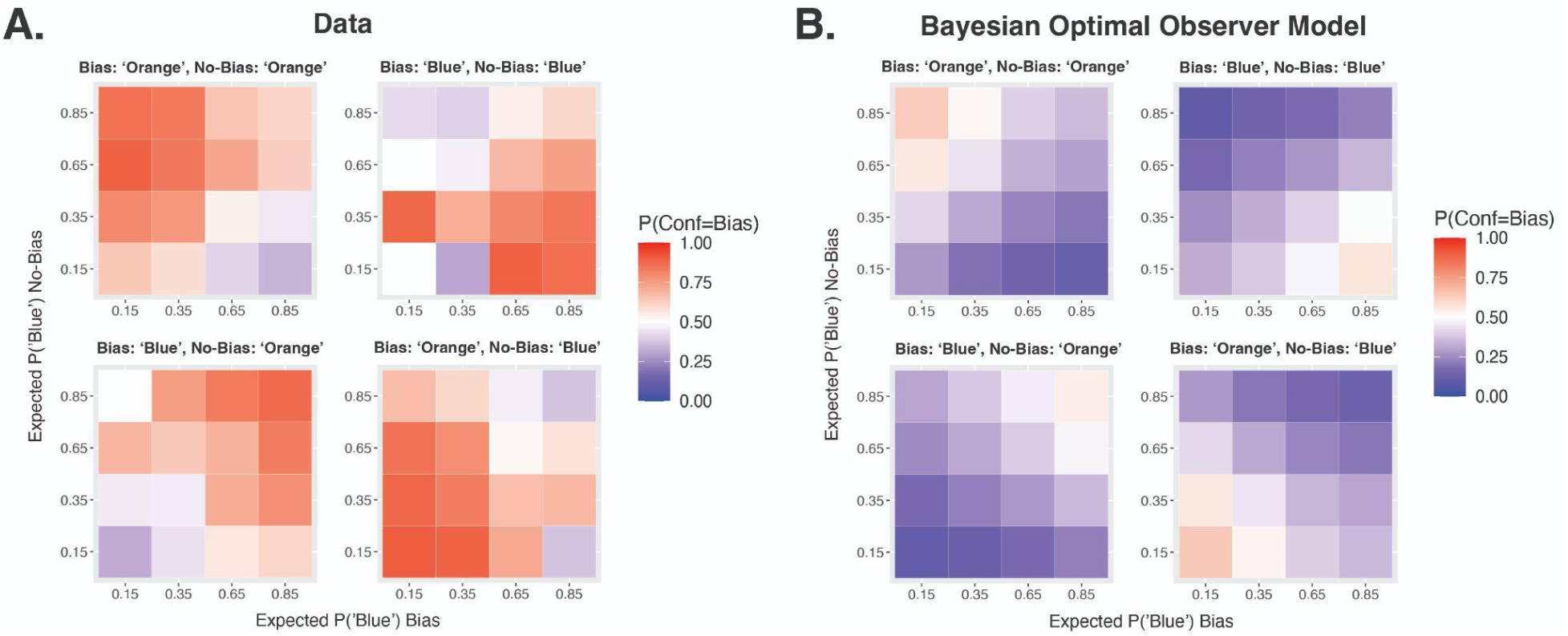
Confidence results and predictions. **A. Confidence forced-choices.** In two consecutive intervals, a Bias and a No-Bias stimulus were presented (or in the reverse order). Because confidence refers to an estimate that a perceptual decision is correct, we consider separately each possible combination of orange/blue responses across the two intervals (thus 4 panels for all the possible combinations). Within a set of responses, each grid cell captures a combination of stimulus settings across the two conditions, with the Bias condition on the x axis and No-Bias condition on the y axis. The stimulus settings are defined by the expected probability of choosing blue (Expected P(‘Blue’)) from the ASA staircase procedure, or in other words, the targeted choice rate. For example, an Expected P(‘Blue’) of 0.65 refers to the stimulus intensity that was chosen from the staircase to target a blue choice rate of 0.65 in that condition. The color code indicates the measured probability of a confidence choice favoring the interval containing the Bias condition. The red dominance overall suggests that participants are more often most confident in the Bias condition interval. **B. Confidence predictions from Bayesian Optimal Observer model.** Confidence forced-choice predictions simulated from the Bayesian Optimal Observer model in the same format as in panel A. The blue dominance overall suggests that this model predicts more confidence choices favoring the No-Bias condition, unlike what is seen in the data.

The raw confidence choices are in striking contrast to the predicted confidence choice rates of the Bayesian optimal observer model, shown in Fig. 4B. The expected rates of confidence choices favoring a condition depend not only on stimulus intensity and response accuracy, but on the relative strengths of the *posteriors* in each condition, and therefore on the use of the prior. In the Bayesian model, because of the shape of the Gaussian mixture priors, the distribution of posteriors across different samples of the stimulus (or in other words across trials) is quite different between the conditions (Fig. 3B). In the No-Bias condition, the maxima of these posteriors are pulled towards the peaks of the prior at either + or −90°, so a large proportion of the area is distributed relatively far from the sensory reference on the chosen side (Fig. 3B). Having a higher proportion of the area on the chosen side like this leads to higher confidence, and therefore the Bayesian optimal observer will on average have higher confidence in the No-Bias condition despite matched first-order choice rates. This pattern is shown clearly in the model simulations of the optimal Bayesian confidence observer given the first-order decision results (Fig. 4B). This suggests that, if our participants used the information from the prior in a Bayesian optimal manner for their confidence, there would have been a confidence bias favoring the No-Bias condition, which is the opposite of what is seen in the data (Fig. 4A). So, we next explore some alternative confidence models that might better explain the patterns found.

### Distance-to-Criterion Confidence Model

In the Bayesian Optimal confidence model, confidence is based on the area under the full Bayesian posterior. However, another possibility for computing confidence is to base it off of a single sample from the posterior, in which case the brain may not need knowledge of the full distributional properties (see, e.g., (36)). In this model, confidence is proportional to the distance between the posterior sample and the sensory reference, or ‘criterion’ (Fig. 5A). Samples further from the criterion very clearly support that choice and lead to higher confidence, whereas samples close to the criterion indicate uncertain choices and lead to lower confidence. This confidence model has no free parameters beyond the first-order sensitivity parameters fit to perceptual decisions and used in all models. We simulated confidence forced-choices from it to investigate whether it could better account for the data. This revealed even more extreme predictions of a confidence bias favoring the No-Bias condition than in the Bayesian Optimal model (Fig. 4B), suggesting this model could only poorly capture the confidence data.

**Figure 5.**
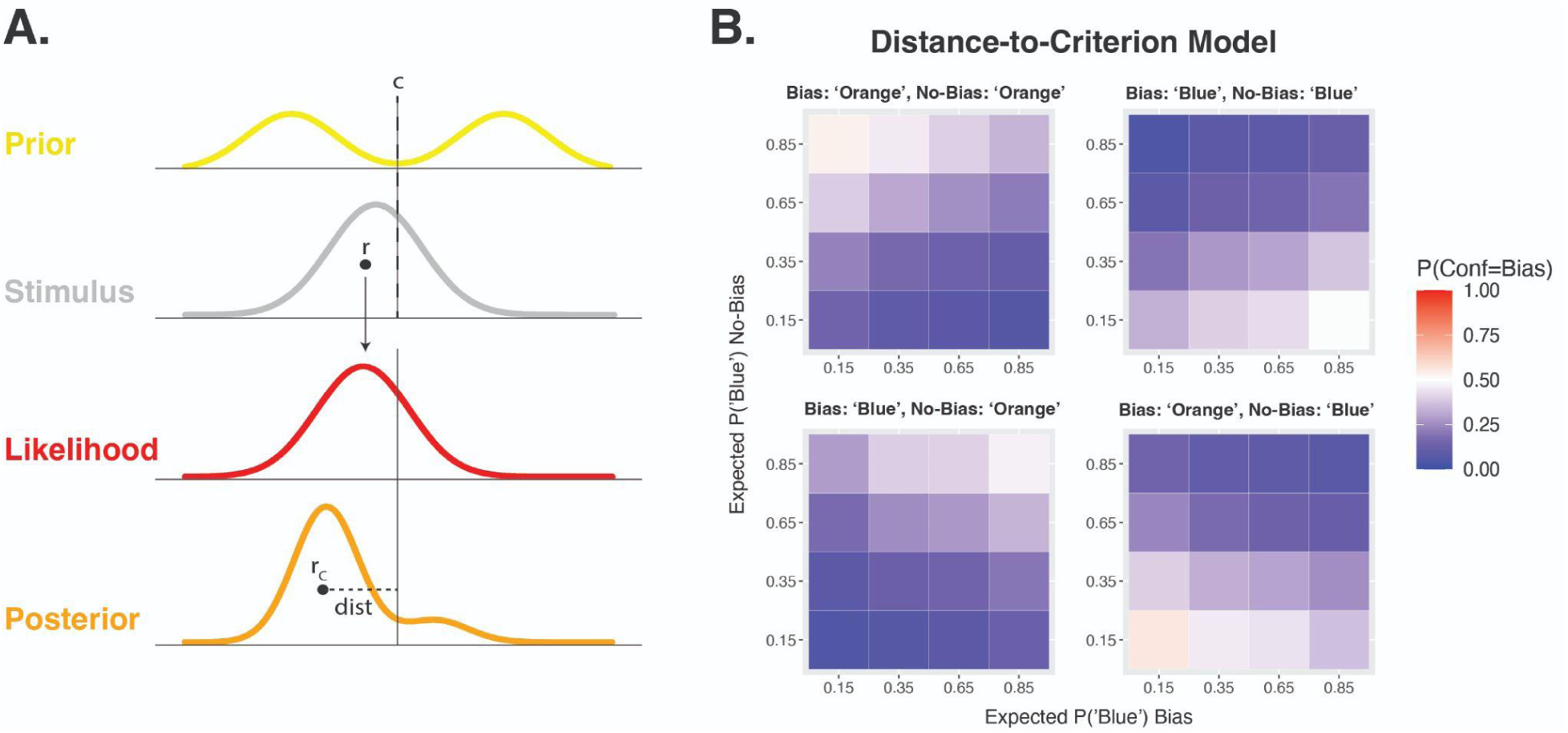
Distance-to-Criterion Confidence Model. **A. Sketch of Distance-to-Criterion model.** In the Distance-to-Criterion model, the prior, likelihood, and resulting posteriors are the same as in the Bayesian Optimal Observer model. Motion-direction decisions also work identically. However, at the confidence level, instead of computing the full Bayesian posterior probability of being correct, this model bases confidence on the absolute distance, *dist*, between a single sample from the posterior, *r_c_*, and the reference, or criterion (*c*). Larger distances lead to higher confidence and smaller distances lead to lower confidence. **B. Confidence predictions from Distance-to-Criterion model.** Confidence forced-choice predictions simulated from the Distance-to-Criterion model in the same format as in Figure 4A. Like in the Bayesian Optimal Observer model, the blue dominance overall suggests that this model predicts more confidence choices favoring the No-Bias condition, unlike what is seen in the data.

### Weighted Prior Confidence Model

The Bayesian Optimal and Distance-to-Criterion models both base the confidence on the same posterior as the first-order decisions, and fail to capture the confidence bias seen in the data favoring the Bias condition. It is possible that the confidence data can be better explained by using a different weighting of the prior relative to likelihood in computing confidence, similar to the pattern found in previous work (18). In the Weighted Prior (WP) model, the variance of the likelihood is scaled by a multiplicative factor *w* during the confidence computation to capture a relative over- or underestimation of the likelihood precision and hence an over- or underweighting of the likelihood information relative to the prior (Fig. 6A). When *w*=1, the likelihood variance is correctly estimated, the relative weighting is optimal, and this model is identical to the Bayesian Optimal confidence model. When *w*>1, the likelihood variance is overestimated and therefore it has a weaker effect on the computation of confidence, and the prior is relatively overweighted in confidence. When *w*<1, the likelihood variance is underestimated and therefore it has a stronger effect and the prior is relatively underweighted in confidence. In essence, this model is a naive implementation of the confidence bias parameter in the Confidence-Forced-Choice model (33) where large values of *w* correspond to small values of *β*. We fit this model with its free parameter *w* to the group data using a maximum likelihood estimation approach. This revealed a best-fitting weighting parameter of *w*=1.54, indicating that the prior is overweighted in confidence. This is in agreement with the results of previous work which also found an overweighting of prior information at the confidence level using a similar model implementation (18).

**Figure 6.**
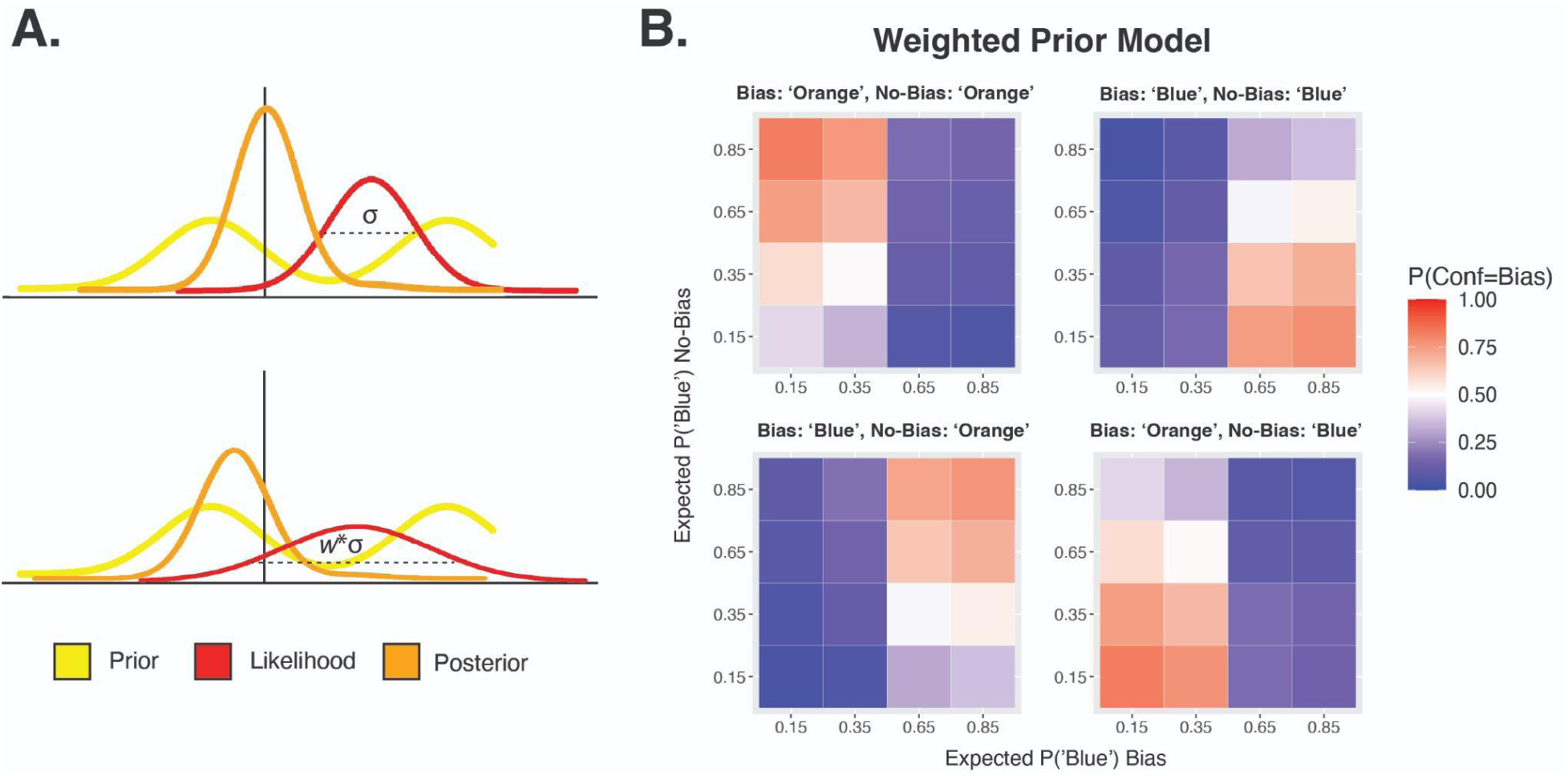
Weighted Prior Confidence Model. **A. Sketch of Weighted Prior model.** In the Weighted Prior model, confidence is computed the same way as in the Bayesian Optimal Observer model, by taking the area under the posterior distribution on the chosen side. However, in forming this posterior, participants over- or underestimate the precision of the likelihood relative to the prior, leading to the prior being relatively over- or underweighted in the confidence computation. The over- or underestimation of the likelihood precision is captured by a scaling term *w* that acts as a multiplicative factor on the variance of the likelihood. When this value is 1, the variance is correctly estimated and this model is the same as the Bayesian Optimal Observer model. When *w*<1, the prior is relatively underweighted and when *w*>1 the prior is relatively overweighted in confidence. **B. Confidence predictions from fit Weighted Prior model.** Confidence forced-choice predictions simulated from the fit Weighted Prior model, with the best-fitting *w*=1.54, in the same format as in Figure 4A. More red in these simulations compared to the Bayesian Optimal Observer model (Fig. 4B) suggests that overweighting prior information can lead to more confidence decisions favoring the Bias condition, closer to what is seen in the data (Fig. 4A).

Simulations from this fit model are shown in Fig. 6B. These demonstrate the ability of this model to better capture the confidence choices favoring the Bias condition than the Bayesian Optimal and Distance-to-Criterion models. However, this model predicts confidence to be particularly high in the Bias condition when the first-order response is in agreement with the prior, and to be particularly low when the first-order response goes against the prior. This leads to the prediction that confidence choices favoring the Bias condition will occur when the Bias condition response was in agreement with the bias direction, and confidence choices favoring the No-Bias condition will occur when the Bias condition response went against the bias direction (Fig. 6B). As *w* gets higher and the prior is weighted more strongly, this pattern gets even more extreme (Fig. S3). This model still cannot qualitatively capture the confidence choice patterns seen in the data, which do not follow this split.

### Prior-Congruent Evidence (PCE) Confidence Model

From the previous models, it seems that confidence forced-choices are biased towards the condition that is more driven by the prior, but not simply by overweighting the prior (or downweighting the likelihood) in the integration to form the posterior. An alternative scenario is that the information from the prior forms its own cue to confidence, separately from the Bayesian posterior evidence used in the perceptual decisions. The Prior-Congruent Evidence confidence model suggests that confidence is based solely on the amount of prior-congruent evidence (PCE) available in the line-motion stimulus, and not on the posterior evidence or the orange/blue decision made, as in the previous models. The prior-congruent evidence refers to the component of the stimulus motion speed that is projected on the axis orthogonal to the lines’ orientation. Confidence here was proportional to the length of this vector component, as shown in Fig. 7A, with no additional free parameters for confidence in this model. There tended to be more of this prior-congruent evidence in the Bias condition, since line motion directions were closer to the preferred orthogonal direction, and hence simulations from this model reveal a confidence bias favoring the Bias condition (Fig. 7B), in agreement with the confidence forced-choice results. However, the predicted confidence bias from this model is stronger than what we see in the data.

**Figure 7.**
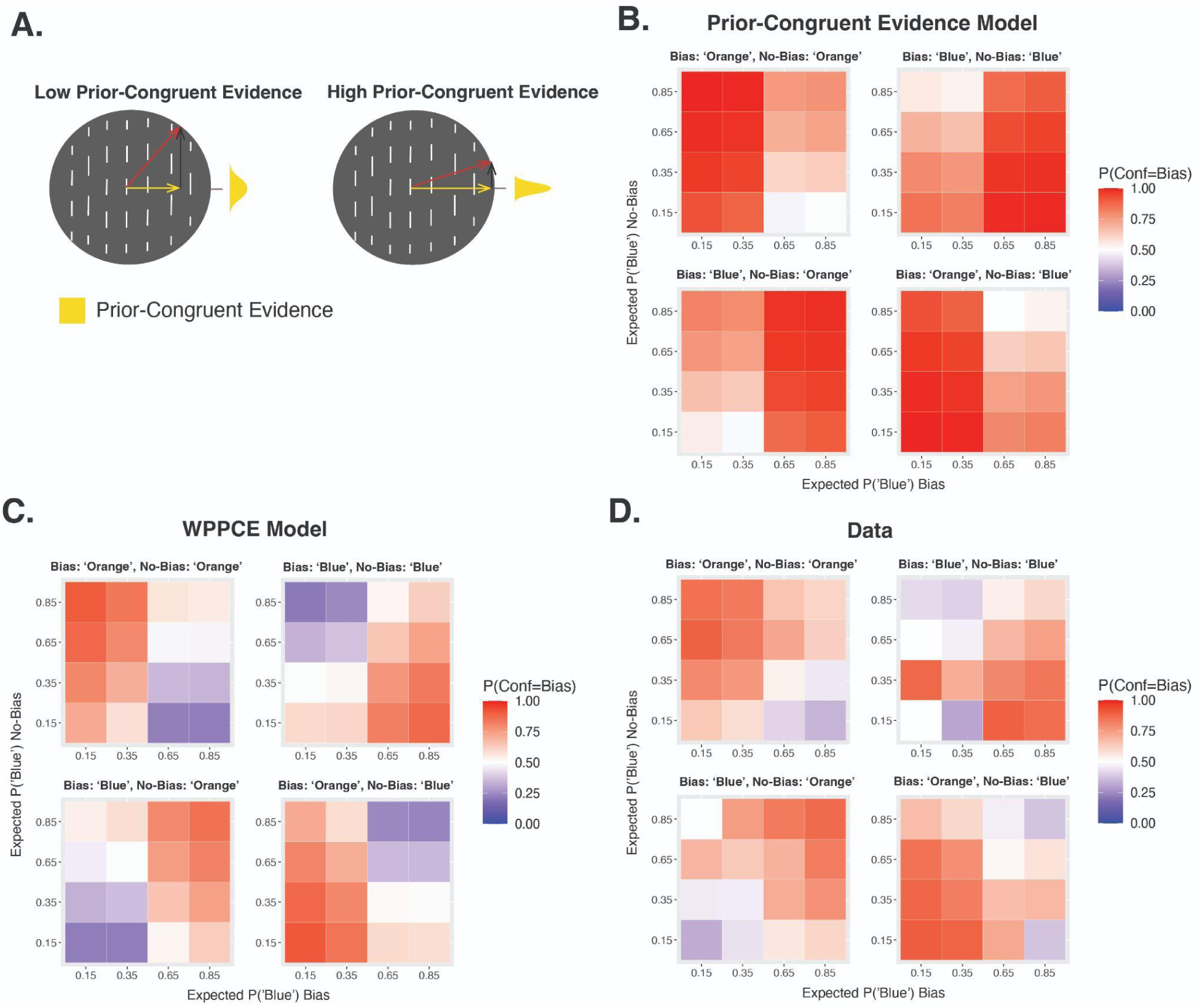
Prior-Congruent Evidence Models. **A. Sketch of prior-congruent evidence.** In the line-motion stimuli, the evidence that is congruent with the prior is the component of the motion that is in the preferred direction, orthogonal to the lines’ orientation, shown as the yellow vector component. The true motion direction (likelihood) is shown in red, and the prior-incongruent component, parallel to the lines’ orientation, is shown in black. Stimuli with high PCE have a motion direction that is closer to the preferred one, and therefore a longer vector component along that dimension. In the PCE confidence model, confidence was directly proportional to this PCE, and was not based on the posterior evidence or on the orange/blue decision made. **B. Confidence predictions from Prior-Congruent Evidence (PCE) model.** Confidence forced-choice predictions simulated from the PCE model in the same format as in Figure 4A. The red dominance overall suggests that this model predicts more confidence choices favoring the Bias condition, in line with, though more extreme than, what is seen in the data. **C. Confidence predictions from Weighted Posterior and Prior-Congruent Evidence (WPPCE) model.** Confidence forced-choice predictions simulated from the WPPCE model with best-fitting ɑ=0.34. In the WPPCE model, confidence is based on a weighted combination of the PCE, as shown in (A), as well as the Bayesian Optimal posterior evidence, as shown in Fig. 4B. **D. Reminder of confidence forced-choice data.** Fig. 4A is reproduced here for ease of comparison to the WPPCE model..

### Weighted Posterior and Prior-Congruent Evidence (WPPCE) Confidence Model

The PCE model reveals that basing confidence on the prior-congruent evidence in the stimulus leads to a confidence bias favoring the Bias condition. However, when confidence is based solely on that PCE, the bias effect is stronger than what we see in the data. Hence, the last model we consider is such that this PCE acts as an additional confidence cue, but not the sole information on which confidence choices are based. In the WPPCE model, confidence is based on a weighted combination of the Bayesian posterior evidence (BPE), that also drives the first-order decisions, as well as the degree of prior-congruent evidence (PCE). The influence of each of these two cues is determined by the weighting parameter, ɑ, such that confidence is proportional to the sum of (1-ɑ)*BPE and ɑ*PCE. Higher ɑ values then lead to more contribution from the PCE. When ɑ=1, the PCE is the only cue to confidence and this is identical to the PCE model, and when ɑ=0, the BPE is the only cue to confidence and this is identical to the Bayesian Optimal Observer model. We fit this free weighting parameter to the group data using a maximum likelihood estimation approach, and found that a value of ɑ=0.34 could best explain the data. Because the BPE and PCE use different units, it is difficult to interpret the raw ɑ value. However, simulations from this fit model are shown in Fig. 7C, and can capture the data well qualitatively, showing a similar confidence bias for the Bias condition that is not solely split based on whether the response was prior-congruent (Fig. 7D).

### Model Comparison

To compare the models quantitatively, we computed the AIC for each, therefore also penalizing the models with more free parameters. This revealed an AIC_Bayesian Optimal_ = 30763.85, AIC_Distance-to-Criterion_ = 38770.61, AIC_WP_ = 29838.48, AIC_PCE_ = 31676.98, and AIC_WPPCE_ = 25851.11 (Fig. 8A). The best model to explain the group confidence forced-choice data is the WPPCE confidence model (Fig. 8B), in which confidence choices are based not only on the posterior evidence that formed the first-order decision, but additionally on the prior-congruent evidence as a separate cue. We additionally fit and compared the five models for each individual participant. This revealed the WPPCE confidence model to be the winning model in 23 out of the 24 participants, with one participant for whom the PCE model had the lowest AIC, although this was inconclusive in comparison to the WPPCE (Fig. 8C). Taken together, these results suggest that, relative to their first-order behavior, participants have a confidence bias that favors the condition in which the slow-motion prior more strongly influences decisions. Further, this bias can best be explained by a model in which participants use the prior-congruent information as an additional cue to inform their confidence, in combination with the Bayesian posterior evidence used for the motion-direction decisions.

**Figure 8.**
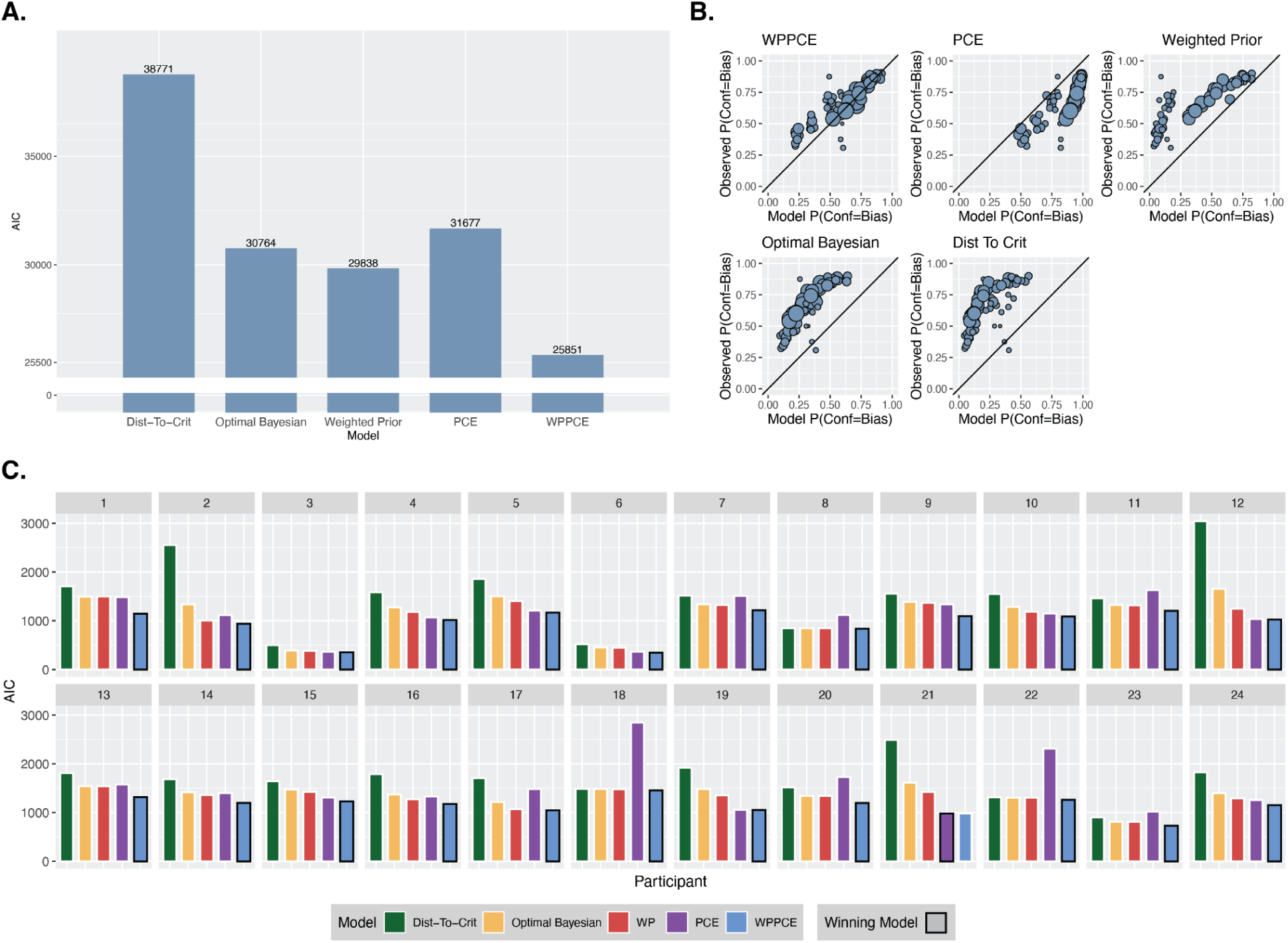
Model Comparison. **A. AIC group results.** AIC comparison of models fit to the pooled group data. **B. Model goodness of fit.** Predicted against observed confidence choice rates from each of the models. Each point reflects a pair of stimulus settings with one interval in each condition and a given set of orange/blue responses, and the size of the point reflects the number of trials. The closer the points are to the x=y line, the better the model is at modeling confidence choice rates. Confidence choice rates reflect the probability of choosing the Bias condition as the more confident interval. So, points above the identity line reflect the model underestimating the frequency of confidence choices for the Bias condition, and points below the identity line reflect the model overestimating the frequency of confidence choices for the Bias condition. **C. Participant-wise AIC results.** AIC comparison of models fit to each individual participant. The winning model for each participant is shown outlined in black. This was the WPPCE model for all except one participant.

## Discussion

Previous work has revealed dissociations between the impact that priors have on confidence versus first-order decisions, suggesting a particularly strong role of priors in shaping confidence (18). However, this previous work examined high-level priors that are induced flexibly in a task context, and may not be generalizable to perceptual priors that are formed naturally across the lifetime. Here, we investigated the role of priors in perceptual decisions and confidence using such a long-term perceptual prior, namely, the slow-motion prior. Using moving line stimuli, we created two conditions: one in which motion direction decision performance was strongly biased by the slow-motion prior (Bias condition), and another in which no bias was induced and performance was driven by the stimulus information (No-Bias condition). We found that, even when any differences in first-order decision rates between conditions were accounted for, there was a confidence bias towards the Bias condition. This suggests confidence to favor the condition with more influence from the prior on decisions, rather than the condition with a more informative stimulus. By comparing computational models, we revealed this effect to be best explained by confidence choices using the prior-congruent evidence as an additional cue, over and above the posterior evidence that is used in first-order decisions.

In line with findings using high-level, flexibly induced priors (18), the results here reveal that long-term perceptual priors also have a different influence on perceptual performance and confidence, with prior-congruent information having a particularly strong impact at the metacognitive level. This underlines a striking consistency of the observed confidence-performance dissociation across different types of priors. Because priors of these different categories are often argued to act and even be implemented differently (21,23), finding this dissociation to hold in the case of long-term priors is enlightening about its nature. Based on work using instructed probabilistic priors, it remained unclear whether their disproportionate impact on confidence was due to the abstract probabilistic information being very unlike (and therefore difficult to integrate with) perceptual information, or due to it acting at a late processing stage. However, long-term priors like the slow-motion prior are assumed to act directly at a perceptual level (26,27,32). Therefore, the results here suggest that this pattern cannot be easily explained by high order, post-perceptual information acting to a greater extent on confidence. Another key difference between the high-level prior manipulations used previously and the low-level one used here is that participants were not aware of the influence of the slow-motion prior. This prior was naturally present in participants, not temporarily manipulated in the lab. Further, the stimuli were designed such that participants would not even notice a decision bias forming from the prior, since it biased their decisions evenly towards blue and orange response sectors. This rules out the possibility that the strong effect of priors on confidence can be explained by participants realizing that they had this prior information and incorporating it post-decisionally, or by a confidence bias induced by demand characteristics. The use of the confidence forced-choice paradigm further eliminates this possibility, reducing the chance of any report- or scale-based confidence biases (33).

Here we find confidence to be biased towards information that is consistent with one’s prior beliefs. This relates to other confirmatory confidence biases in the literature, such as the positive evidence bias in which confidence is biased towards information that confirms one’s decision (37,38). Similarly to the positive evidence bias, we also find this to occur at a low level, and outside of the awareness of participants. Both of these confirmatory confidence biases may be adaptive for avoiding cognitive dissonance, and driving self-consistency (39,40). Also, although confirming prior-congruent information may seem maladaptive for maintaining good predictive models, having a positive epistemic feeling that reinforces prior-congruent information might actually help drive us to minimize prediction errors by encouraging us to seek information that will be in agreement with expectations (1,41). The findings here may also relate to another confidence-performance dissociation that has been discussed in the literature, in which confidence is influenced by the visibility of a stimulus, over and above the total decision evidence (42,43). That work modeled confidence as a weighted combination of these two factors. In parallel with this, the prior-congruent evidence in the WPPCE model proposed here could reflect a visibility-like cue to confidence, if the prior-congruent information increases perceptual strength. This is in agreement with work finding that prior-congruent stimuli enter conscious awareness earlier, and at lower thresholds (44–49). The confidence forced-choice task has a benefit of reducing confounding confidence biases, but it also limits our insight into the subjective quality of the effect. Many different perceptual features have been suggested to have the potential to impact confidence, despite being decision-irrelevant (50). Future work could investigate further in which way information in the preferred orthogonal motion direction impacts the phenomenal experience of the stimuli, which may clarify how the findings here relate to other confidence biases.

### Limitations

Although this work extends previous findings to what is often considered a different category of prior, it is still only one example of a long-term prior, and it is possible that others act in different ways. In line with this possibility, recent work examining confidence under the influence of natural image statistics found somewhat different effects of an orientation prior versus a lighting prior on confidence (19). It would therefore be valuable to find ways to investigate other examples. Also, although we see a similar confidence-performance dissociation with a long-term prior as was previously found with high-level priors, we cannot draw any conclusions about a shared mechanism of action. Future studies testing for correlations between the strengths of these dissociations across different priors, as well as neuroimaging work exploring this effect in terms of neural implementation would be useful to clarify this. Finally, while the conditions we created and models we tested allowed us to quantitatively compare prior-use in decisions and confidence, they also came with some limitations. Our use of a binary discrimination task for the perceptual judgments allowed us to build the conditions such that the prior only influenced decisions in the Bias condition. However, there could still be residual effects of the prior on perception in the No-Bias condition that do not impact performance, which could be explored in future work using a reproduction task. We also note that here we chose to model the prior in terms of the directional bias that is created in these stimuli, rather than modeling the prior over speed. This was for computational tractability of the confidence models as well as simplicity of interpretation, since the directional prior reflects the focus of the motion direction task better. Further, the effect of the slow speed prior on the perceived motion direction bias has been explored extensively in previous work (26,27,32,51) and was not central to our research questions, which only required that we could well capture the resulting bias effect in decisions and confidence. Still, it could be interesting for future work that is more focussed on understanding the nature of the slow speed prior to explore these questions in a speed estimation task.

## Conclusion

While there is evidence from previous work that priors impact first-order decisions and confidence in different ways, it had previously remained unclear whether this would generalize to long-term perceptual priors. Here, we found a similar dissociation in the effect of the slow-motion prior on first-versus second-order processing, with the prior-congruent evidence more strongly impacting confidence. This suggests a confirmatory confidence bias favoring evidence congruent with priors, which occurs implicitly and from low-level priors that are naturally formed in participants. It is therefore important to account for this effect in order to model and understand confidence across the variety of naturalistic situations in which priors influence our processing.

## Materials and Methods

The experiment was pre-registered (https://osf.io/3uc4k), and we respected the pre-registered plan unless stated otherwise.

### Participants

We pre-registered that we would test 25 participants, ensuring at least 20 clean datasets that met the prespecified inclusion criteria, which most importantly included that they showed the basic effects of the prior – that the choice rates were biased towards the orthogonal motion direction in the Bias condition. We tested 25 participants and later excluded the data from one of them because they did not show the basic effect of the prior, leaving data from 24 participants (7 male, 17 female) included in our analyses. Participants were between 21 and 39 years of age (M=24.76, SD=4.28), reported to have normal or corrected-to-normal vision and were fluent in English. Participants were compensated with 10€ per hour and gave signed, informed consent before starting the experiment. The local ethics committee (Comité d’Ethique pour les Recherches en Santé) approved the study, which conformed to the Declaration of Helsinki.

### Setup

The experiment was programmed using MATLAB (52) and Psychtoolbox-3 (53–55). Participants sat in a dark room with their head on a chin and forehead rest placed 57 cm away from a CRT monitor (Vision Mater Pro 454) with a display resolution of 1,920 x 1,200 (refresh rate=60Hz).

### Procedure

#### ASA Procedure

At the beginning of the first session, after receiving verbal instruction about the stimuli, participants completed a short stimulus training consisting of 80 trials across five mini-blocks (16 trials each) in which the line motion stimulus was shown and they input their answer about whether the lines were moving towards the orange or blue region using the mouse. For the first mini-block, the true line motion direction was shown. In addition, the lines started at high contrast so that participants could clearly see and understand the stimuli, and got lower contrast in later blocks (Michelson contrasts: 10%, 10%, 3%, 3%, 3%), and their movement was displayed for a longer duration at first and got progressively shorter in each block (durations: 1000ms, 266ms, 133ms, 133ms, 133ms). After this training, participants completed a staircasing procedure in order for us to select the appropriate stimulus values in each condition for the full experiment, given their performance and their bias. The No-Bias condition consisted of only one setting, with the reference point exactly 90° away from both preferred, orthogonal line orientation directions. The Bias condition consisted of two settings, to capture the bias effect from the influence of the prior in both directions (a blue-bias and an orange-bias), by setting the preferred, orthogonal line orientation either 35° towards the blue region or towards the orange region from the reference (Fig. 1A). For each of these settings, we ran an accelerated stochastic approximation (ASA) staircasing procedure (56) with 4 staircases, set to converge to two choice probability thresholds of 0.25 and 0.75. We then fit a cumulative normal distribution function (**Φ**) with a free noise (***σ****_asa_*) and bias (***μ****_asa_*) parameter to the data from each of these three separate ASA procedures:

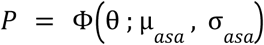

where *P* is the choice probability and ***θ*** is the stimulus value used, which was the angle between the reference and the motion direction (Fig. 1C). These fit psychometric functions were then used to select the stimulus values that targeted probabilities of choosing blue (P(‘Blue’)) of 0.15 and 0.35 when there was an orange bias, and 0.65 and 0.85 when there was a blue bias in the Bias condition, and probabilities of choosing blue of 0.15, 0.35, 0.65, and 0.85 in the No-Bias condition. These stimulus values were used for the main task. This procedure took approximately 10 minutes.

#### Main Task

After completing the ASA procedure, participants received verbal instructions about the structure of the confidence forced-choice task, and completed another short training of 8 trials. Each trial of the main task consisted of two consecutive motion direction decisions followed by a confidence choice. We used this criterion-free confidence forced-choice structure in order to reduce confidence biases that come from criterion placement or particular use of a confidence scale, which can make the interpretation of confidence results difficult (33). The motion direction decisions worked in the same way as in the ASA procedure – participants viewed the set of moving lines at short duration (133ms) and low contrast (3.5% Michelson contrast) and then used the mouse to select whether the motion direction was towards the orange area or blue area by clicking on the corresponding colored quarter-ring around the display circle. In the confidence choice, participants used the left and right arrow keys to indicate whether they were more confident in the first or second interval, respectively (Fig. 1B). We used the chosen stimulus values (θ’s, four per condition) from the ASA procedure to create 28 possible pairs, by using all possible pairings of these eight stimulus values except for ones with the identical stimulus value and condition twice. These 28 pairs formed the possible interval pairs and were counterbalanced across the main task in both possible interval orders. We fixed each block such that half the trials would have the line orientation rotated clockwise from vertical, and half counterclockwise from vertical, and amount of rotation was sampled randomly on each trial from the possible values: 10°, 20°, 30°, 40°, 50°, 60°, 70°, or 80°. We also fixed each block such that half the trials would have orange and half would have blue as the ring color counterclockwise to the reference. After assigning the line orientation and the placement of the blue and orange regions, the rotation of the reference and then line motion direction were set relative to these, and were dictated by the stimulus value pair – the condition dictated the placement of the reference and the θ dictated the line motion direction (Fig. 1A). If the stimulus setting dictated a true line motion direction towards blue, the θ would be set so that the lines moved towards the region colored blue on that trial. In the Bias condition, if the stimulus setting dictated a ‘Blue Bias’, the reference would be set such that the preferred motion direction fell in the region colored blue on that trial. The order of trials was randomized within each block.

Participants completed 24 repetitions of each of the 28 stimulus pairs for a total of 672 trials, including 672 motion direction decisions in each condition (for a total of 1344 motion direction decisions) and 672 confidence choices. These took place across 12 blocks with 56 trials in each block. Four of these blocks were completed in the first session and the remaining eight took place in the second session, with the sessions occurring either on the same day with a minimum of one hour break between them, or on consecutive days. With the time taken for instructions and the ASA procedure in the first session, this resulted in each session taking approximately 1 hour.

#### Stimuli

The line motion stimulus was similar to the stimulus used in work by Sotiropolous et al. (32). It consisted of a matrix of parallel line segments that moved rigidly, all in the same direction, and at a constant velocity. The line movement speed was 3° of visual angle per second and they were shown for a duration of 133ms. The matrix of lines was displayed through a circular mask in the center of the screen, which was 12° of visual angle in diameter. The line segments making up the matrix were 2.86° of visual angle, and were hence much shorter than the circular mask, allowing many endpoints to be visible. This is critical because only the line ends provide the disambiguating information regarding the line velocity. The lines were 3.0 arcmin thick. The background was middle gray with a luminance of 0.5 cd/m^2^ and the lines were lighter gray with a luminance of 0.536 cd/m^2^, giving a Michelson contrast of 3.5%. For the orange/blue motion direction decision, the orange and blue choice regions were displayed by showing a colored half-ring around 180° of the circular mask, with 90° of that being blue and 90° of it being orange. This half-ring was visible for the duration of the line motion in order to avoid it appearing after and masking the stimulus. Participants then reported their decision by clicking the desired region of this half-ring. When participants moved the mouse close enough to one of the choice regions to select it, the orange or blue segment expanded slightly to highlight it, such that participants knew which they were selecting. This was helpful because the mouse cursor was invisible in order to avoid distracting participants during the stimulus display, or having a masking effect if it appeared immediately after. Before each stimulus, a green fixation point in the center of the screen was displayed for 500ms. The half-ring only appeared after this, with the matrix of lines, in order to avoid participants planning a strategy or fixating on the reference prior to the stimulus onset.

### Analysis

As pre-registered, we removed any trials with reaction times that were longer than 8 seconds for any decision, or shorter than 100ms for any motion direction decision. We also removed trials that included a specific stimulus setting for which the basic bias effect was not present.

Our behavioral analysis testing the effect of condition on motion direction decisions was done using a logistic mixed-effects model with the ‘lme4’ package (57) in R (58). The test was two-tailed and used an alpha value of 0.05. We did not assess confidence choices using a regression approach because we prespecified that we would only do this if the first-order choice probabilities were adequately matched, which they were not. Additionally, we realized from further model exploration that the Bayesian optimal observer model does not predict equal confidence given matched first-order choice rates, so this analysis was not the best approach to assess deviations from the Bayesian optimal behavior in any case. Instead, and as planned given that first-order choice rates may not be perfectly matched, to examine confidence bias we used a non-parametric analysis following a similar approach to that in previous work (35). We transformed the choice probabilities to put them in the (-∞,∞) domain by taking their z-score, and then computed the differences between the two conditions for each stimulus pair setting, excluding those with the same condition in both intervals. For each of these z-scored choice probability differences (*cp_diff_*), we then computed the corresponding rate of choosing the Bias condition as the more confident one (*P(“Bias”)*) and fit a cumulative normal distribution function (**Φ**) with a free noise (***σ****_cb_*) and bias (***μ****_cb_*) parameter to this relationship:

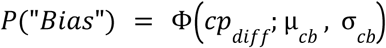

### Modeling

#### Quantifying Confidence Bias

In order to further quantify the confidence bias, before exploring the different process models that might explain it, we fit the *cfc-model* developed in previous work (33). This model includes a confidence bias term, ***β***, which generally captures the propensity to choose one condition as the more confident interval, after accounting for the stimulus strengths, first-order responses in each interval, and a potential interval bias (propensity to prefer a particular interval as the more confident one). In a single condition, confidence bias is captured as a parameter that scales the estimated sensory sensitivity. If this parameter is equal to 1, sensitivity is perfectly estimated. If this parameter is below 1, sensitivity is underestimated indicating underconfidence and if this parameter is above 1, sensitivity is overestimated indicating overconfidence. With two conditions, as in our case, the *cfc-model* fits ***β*** as the ratio of confidence biases between conditions with the No-Bias as the baseline, such that ***β***=1 indicates no confidence bias, ***β***>1 indicates a confidence bias favoring the Bias condition, and ***β***<1 indicates a confidence bias favoring the No-Bias condition. In order to collapse the Bias condition into one psychometric for the *cfc-model*, we projected the orange bias trials onto the blue bias space by taking:

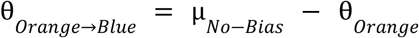

and

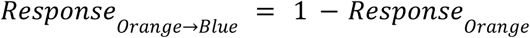

We fit the model to the pooled data, using normalized stimulus intensity values (***θ****_norm_*) that were adjusted based on each participants’ fit sensory noise (***σ****_No-Bias_*) and bias (***μ****_No-Bias_*) from the No-Bias condition:

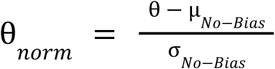

The model freely fit the sensory noise and criteria in each condition, along with the confidence parameters: confidence noise in each condition, confidence boost (reflecting additional information used in confidence) in each condition, interval bias, and the confidence bias parameter, ***β***. We did this for 100 bootstrapped runs of the model in order to extract the confidence intervals surrounding the estimate of ***β***. The other fitted confidence parameter values are reported in Supplementary Information (Table S5). We implemented and fit this model using the previously developed ‘cfc’ software (59). More on the details underlying the model and the other parameters fit can be found in the original work (33).

#### Bayesian Decision Model

To capture the first-order motion direction decisions, we built a Bayesian observer model (Fig. 3A). It has a Gaussian mixture prior distribution of motion directions (in degrees away from the reference) with means at the two preferred, orthogonal motion directions (***μ****_orth1_,* ***μ****_orth2_*),

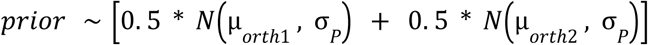

where *σ_P_* is the standard deviation of each Gaussian making up the mixture prior. In the No-Bias condition, ***μ****_orth1_* and ***μ****_orth2_* are at + and −90°. In the Bias condition, ***μ****_orth1_* is 35° away from the reference in the bias direction, and ***μ****_orth2_* is 145° away from the reference in the non-bias direction. The Gaussian likelihood is formed from the stimulus corrupted by sensory noise,

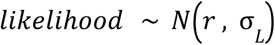

where *σ_L_* is the sensory noise and *r* is the internal signal caused by the stimulus. These combine to form the posterior distribution, and the perceived probability of the motion direction being in the orange (encoded as the negative direction) or blue (encoded as the positive direction) region is computed by the area under this posterior distribution across the entire orange or blue region, normalized:

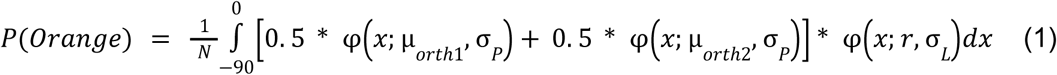

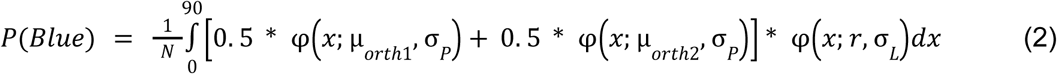

where *N* is equal to the total area under the posterior in the allowed decision region:

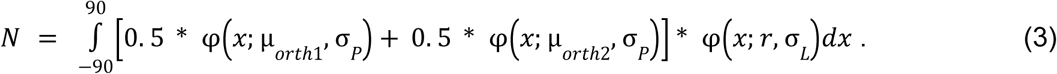

The decision is then based on the max of these two values,

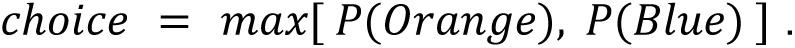

We fit this model to the motion-direction decisions of each individual participant in order to find the *σ_P_* and *σ_L_* values that would best explain their first-order performance. However, in order to deal with the computational demand of this we required a slight simplification. For the Bias condition, we considered the prior to be a single Gaussian centered around ***μ****_orth1_*. Because the Gaussian centered around ***μ****_orth2_* (+-145°) is so far outside of the allowed decision region (from −90° to +90°), this had a negligible impact on the model and made it possible to fit with the amount of data available to us. The ability of the model to still account well for first-order performance can be seen in Fig. 3C. We then used the fit *σ_P_* and *σ_L_* parameters for the rest of the confidence modeling described below.

#### Bayesian Optimal Confidence Model

We then considered the expected confidence patterns of an observer that follows the Bayesian Optimal confidence model. This model is in line with the Bayesian confidence model discussed in the literature in which confidence reflects the perceived posterior probability of being correct about a decision (5–10). In our case, confidence then corresponds to the perceived posterior probability that the orange/blue decision is correct, which is described above in equation (1) if orange is chosen, and (2) if blue is chosen. The confidence forced-choice after both intervals is then based on whichever of these two confidence values from the two intervals is higher:

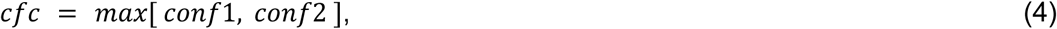

where the *conf* value is the perceived posterior probability correct. This model has no free parameters beyond the first-order sensitivity parameters, *σ_P_* and *σ_L_*.

#### Distance-To-Criterion Confidence Model

It is possible that, instead of computing the area under the posterior distribution as in the Bayesian Optimal confidence model, participants instead just base their confidence on one sample from the posterior. In this model, confidence reflects the distance between a sample from the posterior (*r_c_*) and their criterion (*crit*):

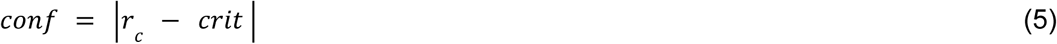

High confidence decisions then occur when the internal posterior sample is far from the criterion, or is more clearly orange or blue. Again, the confidence forced-choice is then based on which of these confidence values is higher, as in equation (4). Like the Bayesian Optimal model, this model has no added free parameters for confidence.

#### Weighted Prior Confidence Model

The Weighted Prior confidence model reflects participants computing confidence as in the Bayesian Optimal confidence model above, but allowing for a different weighting of the prior relative to the likelihood such that the prior may have a stronger or weaker effect in confidence compared to the first-order decisions. The relative over- or underweighting of the prior is implemented by an over- or underestimation of the sensory noise of the likelihood through a scalar weighting parameter *w*, such that the estimated posterior probabilities of each choice at the confidence level are:

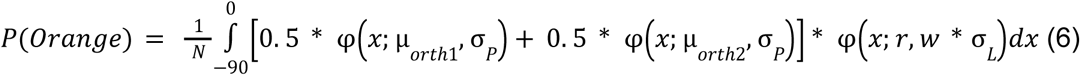

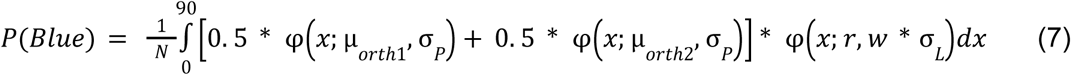

and *N* is:

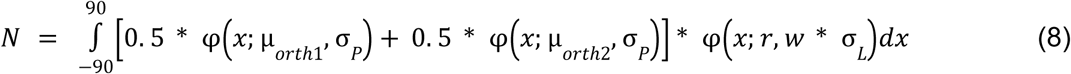

If *w*=1, the sensory precision is correctly estimated in the confidence computation and the prior and likelihood are weighted in the same way as in the first-order decisions. This, additionally, would be identical to the Bayesian Optimal confidence model. If *w*<1, the sensory noise is underestimated, and the prior is underweighted relative to the likelihood. On the contrary, if *w*>1, the sensory noise is overestimated, and the prior is overweighted relative to the likelihood. In this model, the weighting parameter *w* fit as a free parameter. Confidence was allowed to vary between 0 and 1, so there could be cases in which, due to the different prior-weighting at the confidence level, it was below 0.5 and hence did not agree with the perceptual decision. Such cases reflect the possibility of a change of mind following the perceptual decision, with confidence in the originally chosen option then being particularly low.

#### Prior-Congruent Evidence (PCE) Confidence Model

In the PCE confidence model, confidence is not based on the Bayesian posterior distribution but is instead based on the amount of prior-congruent information in the stimulus. For these line motion stimuli, the prior-congruent information refers to the information in the preferred, orthogonal motion direction (Fig. 6A). So, the confidence is proportional to the length of the vector component in this orthogonal direction,

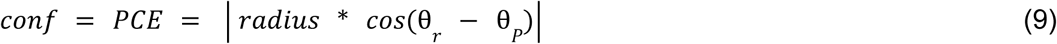

where θ*_r_* is the angle between the reference and the motion direction of the internal signal generated by the stimulus; θ*_P_* is the angle between the reference and the preferred, orthogonal direction; and *radius* is the radius of the stimulus circle, which we set to 1 for simplicity and without loss of generality, since the radius was the same across all trials. The confidence choice is then based on which of these *PCE* values is larger, or which of the two intervals had more prior-congruent evidence, independently of the posterior evidence and the first-order responses.

#### Weighted Posterior and Prior-Congruent Evidence (WPPCE) Confidence Model

The WPPCE confidence model suggests that the prior-congruent evidence described above serves as an additional signal, but not the sole signal, to confidence. In this model, confidence is based on the weighted combination of this *PCE* and the Bayesian posterior evidence (*BPE*), weighted according to a weighting parameter *ɑ*,

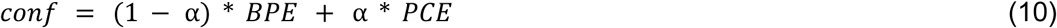

where the *PCE* term is computed as in equation (9), and the *BPE* term is computed as in equation (1) or (2), depending on the response. Larger values of *ɑ* indicate more use of the prior-congruent evidence in confidence, and smaller values of *ɑ* indicate more use of the posterior evidence. When *ɑ*=0, this model is identical to the Bayesian Optimal confidence model and when *ɑ*=1, this model is identical to the PCE confidence model. The weighting term, *ɑ*, was a free parameter.

#### Model-Fitting and Comparison

The confidence models with additional free parameters for confidence – the Weighted Prior model and the WPPCE model – were fit using a maximum likelihood estimation (MLE) approach to find the *w* and *ɑ* parameter values respectively that could best explain the confidence forced-choice data. We used a simulation-based approach to approximate the likelihood, which was analytically intractable. This was also done to compute the likelihood of the models without additional free parameters, to assess model fit. Model fitting was done in R using the ‘stats4’ package. In fitting the Weighted Prior model, we did an initial grid search across values of *w* from 0.1 to 4, in steps of 0.1, and used the *w* value that corresponded with the highest likelihood as the start value of the MLE procedure. This was done to avoid convergence issues. For a model comparison, we compared AIC between the models, computed according to

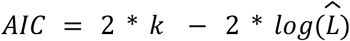

where *k* is the number of free parameters and 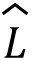 is the maximized value of the likelihood function. We considered a model to be conclusively better if its AIC was lower by at least 2, following convention in the literature. We ensured the models to be distinguishable from one another in a model recovery analysis (Appendix S4; Fig. S4).

## Acknowledgments

MC’s work was funded by the Deutsche Forschungsgemeinschaft (DFG, German Research Foundation)—337619223 / RTG2386. This work was supported by a Freigeist Fellowship from the Volkswagen Foundation (grant number 91620) to EF. EF and PM were supported by a European Commission Doctoral Network grant “CODE” (EC MSCA-101119647). The funders had no role in the conceptualization, design, data collection, analysis, decision to publish, or preparation of the manuscript. Part of this work was presented at VSS 2024.

## Supplementary Information

**Figure S1.**
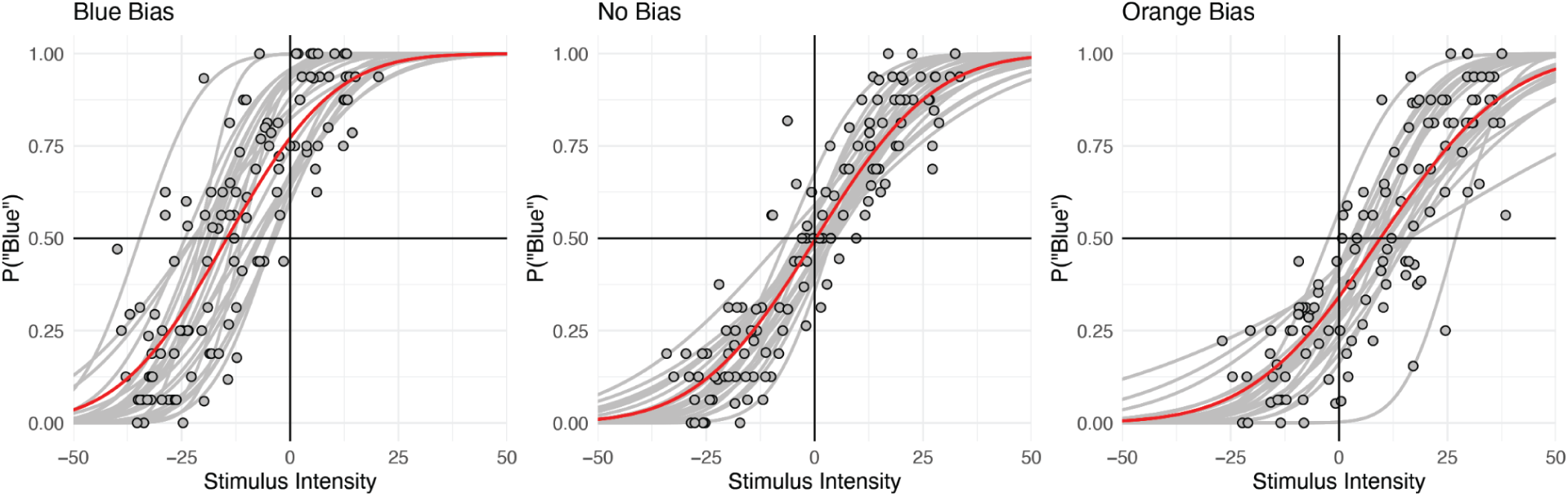
Participant-wise manipulation check. Results from the adaptive staircase (ASA) procedure for each condition across all participants. Each gray point corresponds to the responses for a given stimulus intensity level from one participant. θ values towards the blue region are encoded as positive, and towards the orange region are encoded as negative. Red psychometric functions show the fitted cumulative normal distribution functions to the relationship between θ and the probability of choosing blue from the pooled data. The gray psychometric functions capture the fitted cumulative normal distribution function to data from each individual participant.

**Figure S2.**
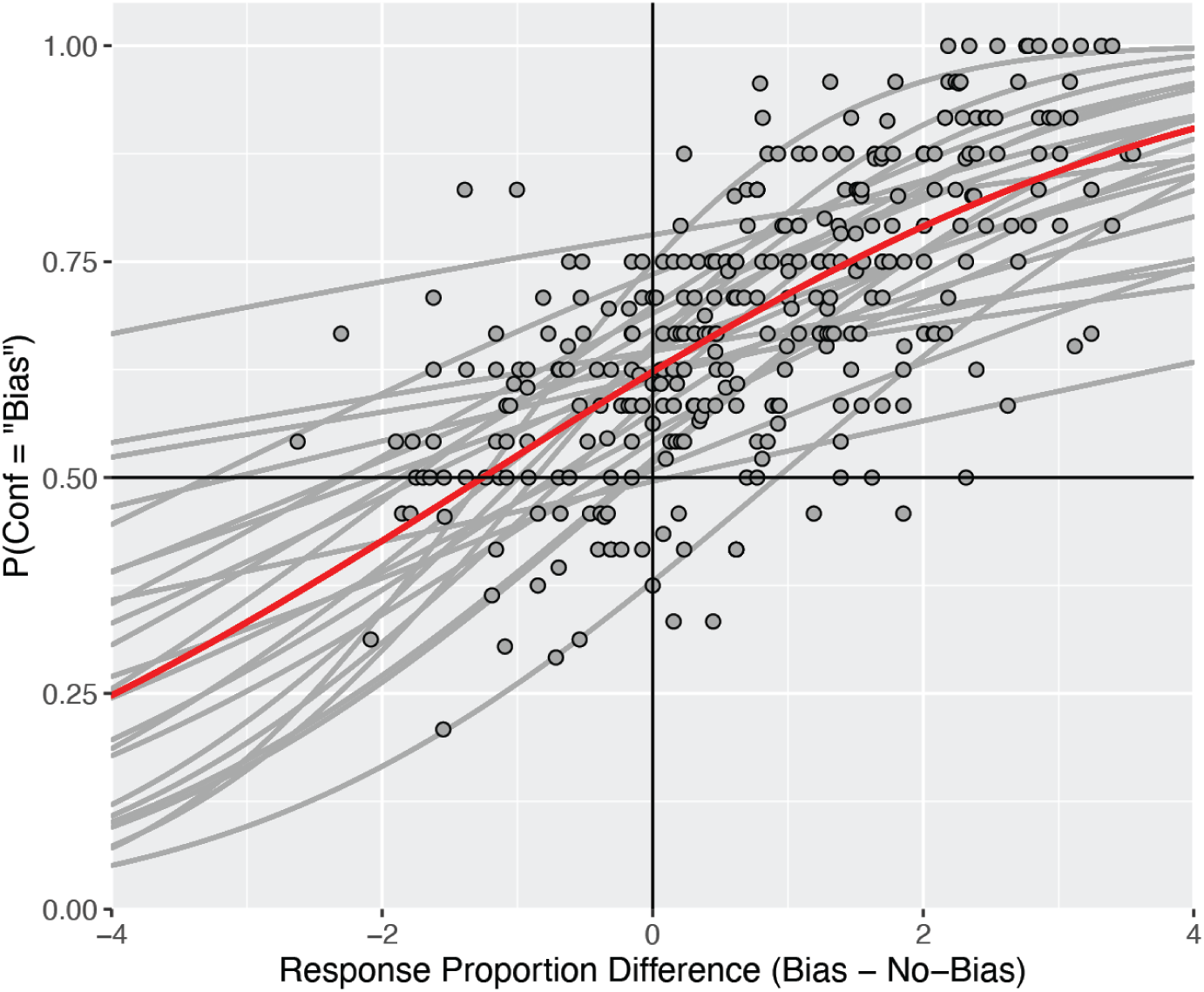
Participant-wise non-parametric analysis of confidence bias. On the x axis is the difference in the z-scored rate of choosing the expected color between the Bias and No-Bias condition. Each gray point represents this difference for one pair of stimulus settings with one interval in each condition, from one participant. On the y axis is the probability of choosing the Bias condition interval as the more confident interval. The red psychometric function captures the fit cumulative normal distribution function to the relationship between this response proportion difference and the confidence choice rates from the pooled data, and the gray psychometric functions capture the fit cumulative normal distribution function to each individual participant.

**Figure S3.**
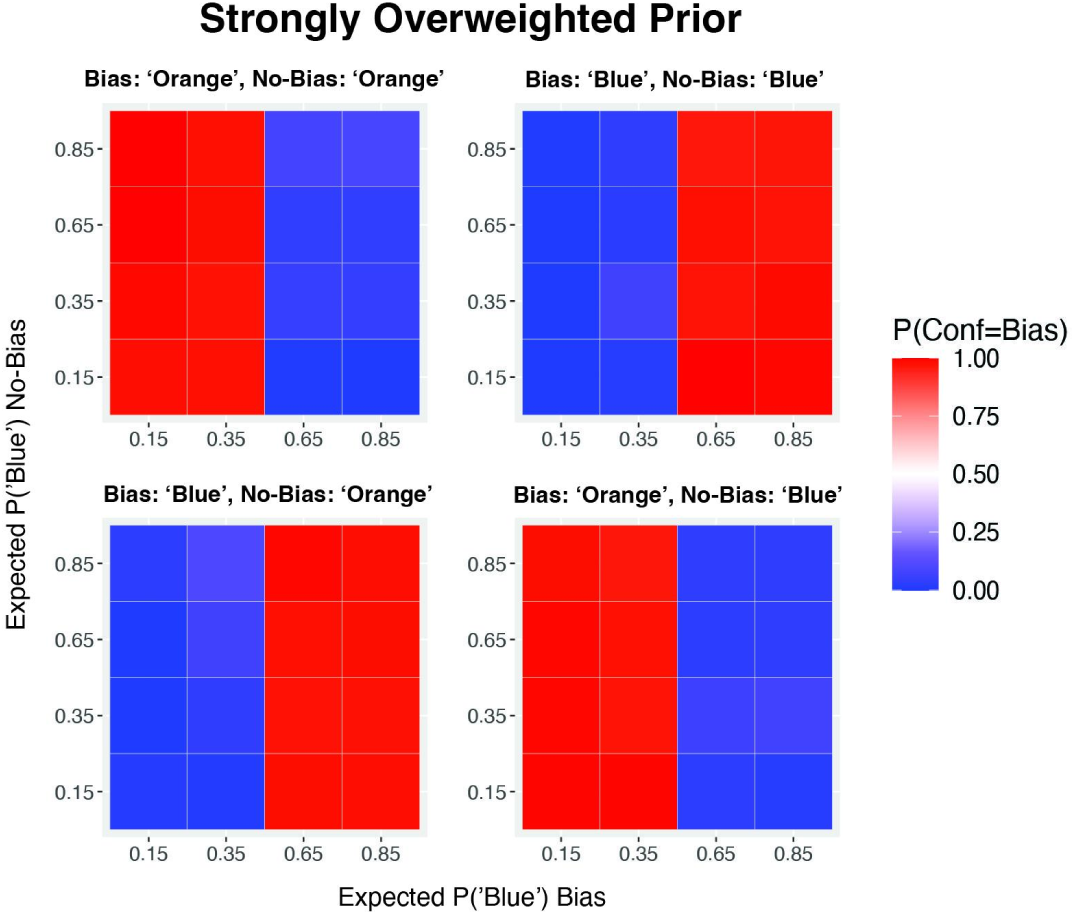
Confidence predictions of extreme prior overweighting. Confidence forced-choice predictions simulated from the Weighted Prior model using an extreme overweighting, *w*=4, for each pair of stimulus settings with one interval in each condition, grouped by possible combination of orange/blue responses. As the overweighting gets extreme, this model predicts the Bias condition to be the confident interval whenever the orange/blue response is in line with the prior, and the No-Bias condition to be the confident interval whenever the orange/blue response goes against the prior.

### Appendix S4. Model recovery analysis

We ran a model recovery analysis in order to ensure that the models were adequately distinguishable in which we simulated data from each model and then checked that the correct model could be recovered. For each of 10 repetitions, we simulated 672 trials of data (equivalent to a single participant) from each model, using representative stimulus values by taking the mean stimulus intensity across participants for each expected performance setting. We also used representative sensory noise values by taking the median fit sensory noise (σ_L_ and σ_P_) parameters across participants, and for the models with additional free parameters for confidence, we used the best fitting values from the group fits: *w*=1.54 for the Weighted Prior (WP) Model and ɑ=0.34 for the WPPCE Model. Models were compared using AIC, with the winning model captured as that with the lowest AIC value. Results of this analysis are shown in

Fig. S4, revealing for each true generative model the proportion of repetitions for which each competing model was recovered as the winning one. This shows that there were only two repetitions in which data were generated from the Bayesian Optimal Observer model and either the WPPCE or WP was recovered as the best model, rather than the correct generative model. This is also not completely unexpected, as both the WPPCE and WP models have certain parameter settings (ɑ=0 and *w*=1, respectively) for which they are indistinguishable from the Bayesian Optimal Observer model. Critically, when data is generated from the WPPCE model with the additional influence of the prior-congruent evidence (ɑ>0), this model is always recovered correctly, suggesting the conclusions from our model comparisons to be reliable.

**Figure S4.**
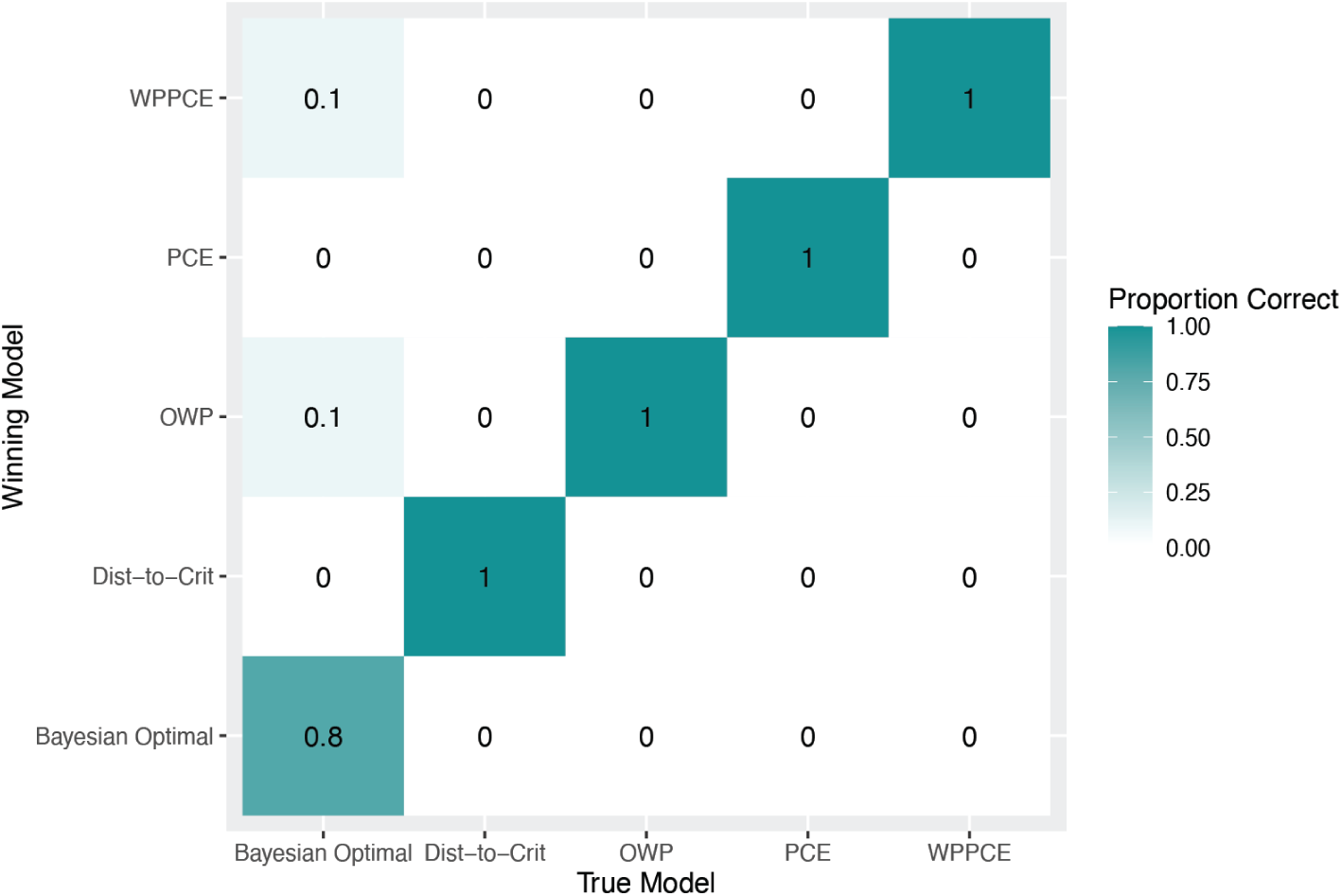
Model recovery analysis. We tested model recovery by simulating data from each model, and then comparing each competing model’s fit to that data using AIC. Columns reflect the true generative model and rows reflect the recovered model. Cell color captures the proportion out of the 10 repetitions for which each competing model (row) was recovered as the winning one, given the true generative model (column). Cells diverging from the diagonal therefore indicate cases for which the incorrect model was recovered as the best fitting one.

### Appendix S5. CFC-model parameters

**Table S5.** Fitted confidence parameters from CFC-model.

| Parameter | Condition | Fitted Value |
| --- | --- | --- |
| Confidence Noise, $\sigma_c$ | Bias | 1.07 |
|  | No-Bias | 0.54 |
| Confidence Boost, $\alpha$ | Bias | 0.83 |
|  | No-Bias | 0.09 |
| Confidence Efficiency, $\eta$ | Bias | 0.78 |
|  | No-Bias | 0.59 |
| Interval Bias, $\gamma$ | All | -0.26 |

## References

1. Friston K, FitzGerald T, Rigoli F, Schwartenbeck P, O’Doherty J, Pezzulo G. Active inference and learning. Neurosci Biobehav Rev. 2016 Sep 1;68:862–79.

2. Kersten D, Mamassian P, Yuille A. Object Perception as Bayesian Inference. Annu Rev Psychol. 2004;55(1):271–304.

3. McNamara JM, Green RF, Olsson O. Bayes’ theorem and its applications in animal behaviour. Oikos. 2006;112(2):243–51.

4. Zellner A. Bayesian and non-Bayesian approaches to statistical inference and decision-making. J Comput Appl Math. 1995 Nov 30;64(1):3–10.

5. Aitchison L, Bang D, Bahrami B, Latham PE. Doubly Bayesian Analysis of Confidence in Perceptual Decision-Making. PLOS Comput Biol. 2015 Oct 30;11(10):e1004519.

6. Fleming SM, Daw ND. Self-Evaluation of Decision-Making: A General Bayesian Framework for Metacognitive Computation. Psychol Rev. 2017 Jan;124(1):91–114.

7. Kepecs A, Mainen ZF. A computational framework for the study of confidence in humans and animals. Philos Trans R Soc B Biol Sci. 2012 May 19;367(1594):1322–37.

8. Meyniel F, Sigman M, Mainen ZF. Confidence as Bayesian Probability: From Neural Origins to Behavior. Neuron. 2015 Oct 7;88(1):78–92.

9. Pouget A, Drugowitsch J, Kepecs A. Confidence and certainty: distinct probabilistic quantities for different goals. Nat Neurosci. 2016 Mar;19(3):366–74.

10. Sanders JI, Hangya B, Kepecs A. Signatures of a Statistical Computation in the Human Sense of Confidence. Neuron. 2016 May 4;90(3):499–506.

11. Sherman MT, Seth AK, Barrett AB, Kanai R. Prior expectations facilitate metacognition for perceptual decision. Conscious Cogn. 2015 Sep 1;35:53–65.

12. Locke SM, Gaffin-Cahn E, Hosseinizaveh N, Mamassian P, Landy MS. Priors and payoffs in confidence judgments. Atten Percept Psychophys. 2020 Aug 1;82(6):3158–75.

13. Sherman MT, Kanai R, Seth AK, VanRullen R. Rhythmic Influence of Top–Down Perceptual Priors in the Phase of Prestimulus Occipital Alpha Oscillations. J Cogn Neurosci. 2016 Sep 1;28(9):1318–30.

14. Schmack K, Bosc M, Ott T, Sturgill JF, Kepecs A. Striatal dopamine mediates hallucination-like perception in mice. Science. 2021 Apr 2;372(6537):eabf4740.

15. Olawole-Scott H, Yon D. Expectations about precision bias metacognition and awareness. J Exp Psychol Gen. 2023 Mar 27;

16. Sherman MT, Seth AK. Effects of expected task difficulty on metacognitive confidence and multitasking [Internet]. PsyArXiv [Preprint]. 2021 [cited 2021 Apr 24]. Available from: https://psyarxiv.com/3gfp2/

17. Van Marcke H, Denmat PL, Verguts T, Desender K. Manipulating Prior Beliefs Causally Induces Under- and Overconfidence. Psychol Sci. 2024 Apr 1;35(4):358–75.

18. Constant M, Pereira M, Faivre N, Filevich E. Prior information differentially affects discrimination decisions and subjective confidence reports. Nat Commun. 2023 Sep 6;14(1):5473.

19. West RK, A-Izzeddin EJ, Sewell DK, Harrison WJ. Priors for natural image statistics inform confidence in perceptual decisions. Conscious Cogn. 2025 Feb 1;128:103818.

20. Mamassian P, Goutcher R. Prior knowledge on the illumination position. Cognition. 2001 Aug;81(1):B1–9.

21. de Lange FP, Heilbron M, Kok P. How Do Expectations Shape Perception? Trends Cogn Sci. 2018 Sep 1;22(9):764–79.

22. Seriès P, Seitz AR. Learning what to expect (in visual perception). Front Hum Neurosci. 2013 Oct 24;7:668.

23. Teufel C, Fletcher PC. Forms of prediction in the nervous system. Nat Rev Neurosci. 2020 Apr;21(4):231–42.

24. Gerardin P, Kourtzi Z, Mamassian P. Prior knowledge of illumination for 3D perception in the human brain. Proc Natl Acad Sci. 2010 Sep 14;107(37):16309–14.

25. Hardstone R, Zhu M, Flinker A, Melloni L, Devore S, Friedman D, et al. Long-term priors influence visual perception through recruitment of long-range feedback. Nat Commun. 2021 Nov 1;12(1):6288.

26. Weiss Y, Simoncelli EP, Adelson EH. Motion illusions as optimal percepts. Nat Neurosci. 2002 Jun;5(6):598–604.

27. Stocker AA, Simoncelli E. Constraining a Bayesian Model of Human Visual Speed Perception. In: Advances in Neural Information Processing Systems [Internet]. MIT Press; 2004 [cited 2024 Apr 15]. Available from: https://proceedings.neurips.cc/paper/2004/hash/852c44ddce7e0c7e4c64d86147300831-Abstract.html

28. Stocker AA, Simoncelli EP. A Bayesian Model of Conditioned Perception. Adv Neural Inf Process Syst. 2007;2007:1409–16.

29. Zhang LQ, Stocker AA. Prior Expectations in Visual Speed Perception Predict Encoding Characteristics of Neurons in Area MT. J Neurosci. 2022 Apr 6;42(14):2951–62.

30. Montagnini A, Mamassian P, Perrinet L, Castet E, Masson GS. Bayesian modeling of dynamic motion integration. J Physiol Paris. 2007;101(1–3):64–77.

31. Gekas N, Mamassian P. Adaptation to one perceived motion direction can generate multiple velocity aftereffects. J Vis. 2021 May 18;21(5):17.

32. Sotiropoulos G, Seitz AR, Seriès P. Changing expectations about speed alters perceived motion direction. Curr Biol CB. 2011 Nov 8;21(21):R883–884.

33. Mamassian P, de Gardelle V. Modeling perceptual confidence and the confidence forced-choice paradigm. Psychol Rev. 2022;129(5):976–98.

34. Mamassian P. Confidence Forced-Choice and Other Metaperceptual Tasks. Perception. 2020 Jun;49(6):616–35.

35. Toscani M, Mamassian P, Valsecchi M. Underconfidence in peripheral vision. J Vis. 2021 Jun 9;21(6):2.

36. Locke SM, Landy MS, Mamassian P. Suprathreshold perceptual decisions constrain models of confidence. PLOS Comput Biol. 2022 Jul 27;18(7):e1010318.

37. Peters MAK, Thesen T, Ko YD, Maniscalco B, Carlson C, Davidson M, et al. Perceptual confidence neglects decision-incongruent evidence in the brain. Nat Hum Behav. 2017 Jul 10;1(7):1–8.

38. Rollwage M, Loosen A, Hauser TU, Moran R, Dolan RJ, Fleming SM. Confidence drives a neural confirmation bias. Nat Commun. 2020 May 26;11(1):2634.

39. Caziot B, Mamassian P. Perceptual confidence judgments reflect self-consistency. J Vis. 2021 Nov 1;21(12):8.

40. Festinger L. A Theory of Cognitive Dissonance. Stanford University Press; 1957. 308 p.

41. Wiese W, Metzinger T. Vanilla PP for Philosophers: A Primer on Predictive Processing. In: Metzinger T, Wiese W, editors. Philosophy and Predictive Processing. 2017.

42. Rausch M, Hellmann S, Zehetleitner M. Confidence in masked orientation judgments is informed by both evidence and visibility. Atten Percept Psychophys. 2018 Jan;80(1):134–54.

43. Rausch M, Zehetleitner M. The folded X-pattern is not necessarily a statistical signature of decision confidence. PLOS Comput Biol. 2019 Oct 21;15:e1007456.

44. Aru J, Rutiku R, Wibral M, Singer W, Melloni L. Early effects of previous experience on conscious perception. Neurosci Conscious. 2016 Jan 1;2016(1):niw004.

45. Eger E, Henson RN, Driver J, Dolan RJ. Mechanisms of top-down facilitation in perception of visual objects studied by FMRI. Cereb Cortex N Y N 1991. 2007 Sep;17(9):2123–33.

46. Esterman M, Yantis S. Perceptual expectation evokes category-selective cortical activity. Cereb Cortex N Y N 1991. 2010 May;20(5):1245–53.

47. Melloni L, Schwiedrzik CM, Müller N, Rodriguez E, Singer W. Expectations change the signatures and timing of electrophysiological correlates of perceptual awareness. J Neurosci Off J Soc Neurosci. 2011 Jan 26;31(4):1386–96.

48. Pinto Y, van Gaal S, de Lange FP, Lamme VAF, Seth AK. Expectations accelerate entry of visual stimuli into awareness. J Vis. 2015 Jun 26;15(8):13.

49. Stein T, Peelen MV. Content-specific expectations enhance stimulus detectability by increasing perceptual sensitivity. J Exp Psychol Gen. 2015;144(6):1089–104.

50. Shekhar M, Rahnev D. Sources of Metacognitive Inefficiency. Trends Cogn Sci. 2021 Jan 1;25(1):12–23.

51. Stocker AA, Simoncelli EP. Noise characteristics and prior expectations in human visual speed perception. Nat Neurosci. 2006 Apr;9(4):578–85.

51. The MathWorks Inc. MATLAB version: 9.9.0 (R2020b) [Internet]. Natick, Massachusetts: The MathWorks Inc.; 2020. Available from: https://www.mathworks.com

52. Brainard DH. The Psychophysics Toolbox. Spat Vis. 1997;10(4):433–6.

53. Kleiner M, Brainard D, Pelli D, Ingling A, Murray R, Broussard C. What’s new in psychtoolbox-3. Perception. 2007;36(14):1–16.

54. Pelli DG. The VideoToolbox software for visual psychophysics: transforming numbers into movies. Spat Vis. 1997;10(4):437–42.

55. Kesten H. Accelerated Stochastic Approximation. Ann Math Stat. 1958;29(1):41–59.

56. Bates D, Mächler M, Bolker B, Walker S. Fitting Linear Mixed-Effects Models Using lme4. J Stat Softw. 2015 Oct 7;67(1):1–48.

57. R Core Team. R: A language and environment for statistical computing. [Internet]. Vienna, Austria: R Foundation for Statistical Computing; 2021. Available from: https://www.R-project.org/

58. Mamassian P. cfc [Internet]. 2023. Available from: 10.5281/zenodo.7791435

